# Advances in Spiral fMRI: A High-resolution Study with Single-shot Acquisition

**DOI:** 10.1101/842179

**Authors:** Lars Kasper, Maria Engel, Jakob Heinzle, Matthias Mueller-Schrader, Nadine N. Graedel, Jonas Reber, Thomas Schmid, Christoph Barmet, Bertram J. Wilm, Klaas Enno Stephan, Klaas P. Pruessmann

## Abstract

Spiral fMRI has been put forward as a viable alternative to rectilinear echo-planar imaging, in particular due to its enhanced average k-space speed and thus high acquisition efficiency. This renders spirals attractive for contemporary fMRI applications that require high spatiotemporal resolution, such as laminar or columnar fMRI. However, in practice, spiral fMRI is typically hampered by its reduced robustness and ensuing blurring artifacts, which arise from imperfections in both static and dynamic magnetic fields.

Recently, these limitations have been overcome by the concerted application of an expanded signal model that accounts for such field imperfections, and its inversion by iterative image reconstruction. In the challenging ultra-high field environment of 7 Tesla, where field inhomogeneity effects are aggravated, both multi-shot and single-shot 2D spiral imaging at sub-millimeter resolution was demonstrated with high depiction quality and anatomical congruency.

In this work, we further these advances towards a time series application of spiral readouts, namely, single-shot spiral BOLD fMRI at 0.8 mm in-plane resolution. We demonstrate that high-resolution spiral fMRI at 7 T is not only feasible, but delivers both excellent image quality, BOLD sensitivity, and spatial specificity of the activation maps, with little artifactual blurring. Furthermore, we show the versatility of the approach with a combined in/out spiral readout at a more typical resolution (1.5 mm), where the high acquisition efficiency allows to acquire two images per shot for improved sensitivity by echo combination.

**Highlights:**

- This work reports the first fMRI study at 7T with high-resolution spiral readout gradient waveforms.
- We achieve spiral fMRI with sub-millimeter resolution (0.8 mm, FOV 230 mm), acquired in a single shot (36 slices in 3.3 s).
- Spiral images exhibit intrinsic geometric congruency to anatomical scans, and spatially specific activation patterns.
- Image reconstruction rests on a signal model expanded by measured trajectories and static field maps, inverted by cg-SENSE.
- We assess generalizability of the approach for spiral in/out readouts, providing two images per shot (1.5 mm resolution).

## 1 Introduction

Functional MRI (fMRI) is presently the most prominent technique to study human brain function non-invasively, owing to its favorable spatiotemporal resolution regime with appealing functional sensitivity. Within this regime, specific research questions require different trade-offs between spatial and temporal resolution. On the one hand, ultra-high spatial resolution fMRI (with sub-millimeter voxel size) successfully targets smaller organizational structures of the brain, such as cortical laminae (Fracasso et al., 2016; Huber et al., 2017a; Kashyap et al., 2018; Kok et al., 2016; Lawrence et al., 2018; Martino et al., 2015; Muckli et al., 2015; Siero et al., 2011) and columns (Cheng et al., 2001; Feinberg et al., 2018; Yacoub et al., 2008). For subcortical sites, due to the limited signal-to-noise ratio (SNR), high-resolution (1-1.5 mm) fMRI is more prevalent (but see (Wang et al., 2020)) to characterize, for example, the superior (Savjani et al., 2018; Singh et al., 2018) and inferior colliculi (De Martino et al., 2013; Sitek et al., 2019), as well as the subthalamic nucleus (de Hollander et al., 2017) and midbrain (D’Ardenne et al., 2008). However, both high and ultra-high spatial resolution fMRI require compromises on field of view (FOV) coverage or temporal bandwidth, i.e., volume repetition time (TR). On the other hand, fast sequences with TRs on the order of 0.5 seconds and below are important for advanced analysis approaches, for example, to adequately sample physiological fluctuations (Lewis et al., 2016; Smith et al., 2013; Uğurbil et al., 2013), at the expense of lowering spatial resolution (2–4 mm).

One means to simultaneously advance the spatial and temporal resolution boundaries of fMRI is to maximize acquisition efficiency, i.e., sampled k-space area (or volume) per unit time. Therefore, fMRI nowadays almost exclusively relies on rectilinear echo-planar imaging (EPI, (Cohen and Schmitt, 2012; Mansfield, 1977; Schmitt et al., 2012)), where acquisition efficiency is favorable due to optimal acceleration and high terminal velocity along the straight k-space lines traversed.

To expand spatiotemporal resolution beyond the capabilities of EPI alone, the main strategy has been parallel imaging acceleration (Griswold et al., 2002; Pruessmann et al., 1999; Sodickson and Manning, 1997), in combination with simultaneous multi-slice or 3D excitation (Breuer et al., 2006; Larkman et al., 2001; Poser et al., 2010; Setsompop et al., 2012). In terms of k-space coverage per unit time, the benefit of parallel imaging lies in expanding the cross section of the k-space neighborhood covered along the readout trajectory (Pruessmann, 2006), i.e., a band in 2D or tube in 3D.

However, another key determinant of acquisition efficiency or speed of coverage is average velocity along the trajectory, i.e., instantaneous gradient strength. On this count, EPI is wasteful because it includes many sharp turns traversed at low speed due to the limited gradient slew rate.

Substantially higher average k-space speed and thus acquisition efficiency for fMRI is achieved with spiral trajectories (Barth et al., 1999; Glover, 2012; Noll et al., 1995), which avoid sharp turns by distributing curvature more evenly (Ahn et al., 1986; Likes, 1981; Meyer et al., 1992). Typically, single-shot variants winding out of k-space center, e.g., on an Archimedean spiral, are prevalent (Glover, 1999; Meyer et al., 1992; Weiger et al., 2002), but different acquisition schemes, such as spiral-in (Börnert et al., 2000) or combined in/out readouts (Glover and Law, 2001; Glover and Thomason, 2004) have been proposed. High-resolution fMRI studies have occasionally employed spirals as well (Jung et al., 2013; Singh et al., 2018), including first applications of laminar fMRI (Ress et al., 2007) and regionally optimized acquisitions, e.g., for the hippocampus (Preston et al., 2010) or superior colliculus (Katyal et al., 2010; Savjani et al., 2018). Common to these approaches is a reduction of acquisition efficiency in favor of robustness by acquiring k-space in multiple shots with shorter spiral readouts.

Despite these efforts, routine use of spiral fMRI has not been established, due to the following three challenges (Block and Frahm, 2005; Börnert et al., 1999): First, spirals are sensitive to imperfect magnetic field dynamics (drifts, eddy currents and other gradient imperfections) which lead to blurring and image distortions. Secondly, non-uniformity of the static B_0_ field, caused by varying susceptibility of the imaged tissues, likewise causes blurring (Bernstein et al., 2004, Chap. 17). Finally, in combination with parallel imaging, spirals pose a somewhat greater reconstruction challenge than Cartesian trajectories (Pruessmann et al., 2001).

Recently, these obstacles have been overcome (Engel et al., 2018; Kasper et al., 2018; Wilm et al., 2017) by (1) employing an expanded signal model that incorporates coil sensitivity encoding as well as independently measured static and dynamic field imperfections (Wilm et al., 2011), and (2) the inversion of this model by an accompanying iterative image reconstruction (Barmet et al., 2005; Man et al., 1997; Pruessmann et al., 2001; Sutton et al., 2003). This approach enabled the use of long spiral readouts (on the order of 50 ms at 7 Tesla), while maintaining high image quality and anatomical fidelity. In particular, such enhanced spiral acquisition efficiency was demonstrated by accomplishing T_2_*-weighted images with a nominal in-plane resolution of 0.8 mm in a single shot. Ultimately, these findings hold promise that spiral fMRI can now indeed profit from the theoretical benefits of enhanced acquisition efficiency to expand the spatiotemporal boundaries of fMRI.

Based on these advances in expanded signal modeling and inversion, in this work, we explore the feasibility and utility of sub-millimeter single-shot spiral fMRI. Specifically, we first assess image quality and temporal stability of fMRI time series obtained with the expanded signal model and algebraic reconstruction. We further evaluate the resulting functional sensitivity and spatial specificity of reference activation patterns, elicited by an established visual quarter-field stimulation paradigm.

Finally, we explore the versatility of the approach with a combined in/out spiral readout at a more typical resolution (1.5 mm). Here, two images per shot can be acquired, translating the high acquisition efficiency of the spiral into enhanced functional sensitivity by echo combination (Glover and Law, 2001; Glover and Thomason, 2004; Law and Glover, 2009).

## 2 Methods

### 2.1 Setup

All data was acquired on a Philips Achieva 7 Tesla MR System (Philips Healthcare, Best, The Netherlands), with a quadrature transmit coil and 32-channel head receive array (Nova Medical, Wilmington, MA, USA).

Concurrent magnetic field monitoring was performed using 16 fluorine-based NMR field probes, which were integrated into the head setup via a laser-sintered nylon frame positioned between transmit and receive coil (Fig. 1 in Engel et al., 2018). Probe data were recorded and preprocessed (filtering, demodulation) on a dedicated acquisition system (Dietrich et al., 2016a). The final extraction of probe phase evolution and projection onto a spherical harmonic basis set (Barmet et al., 2008) was performed on a PC, yielding readout time courses of global phase *k*_0_ and k-space coefficients *k_x_*, *k_y_*, *k_z_* with 1 MHz bandwidth.

**Figure 1.**
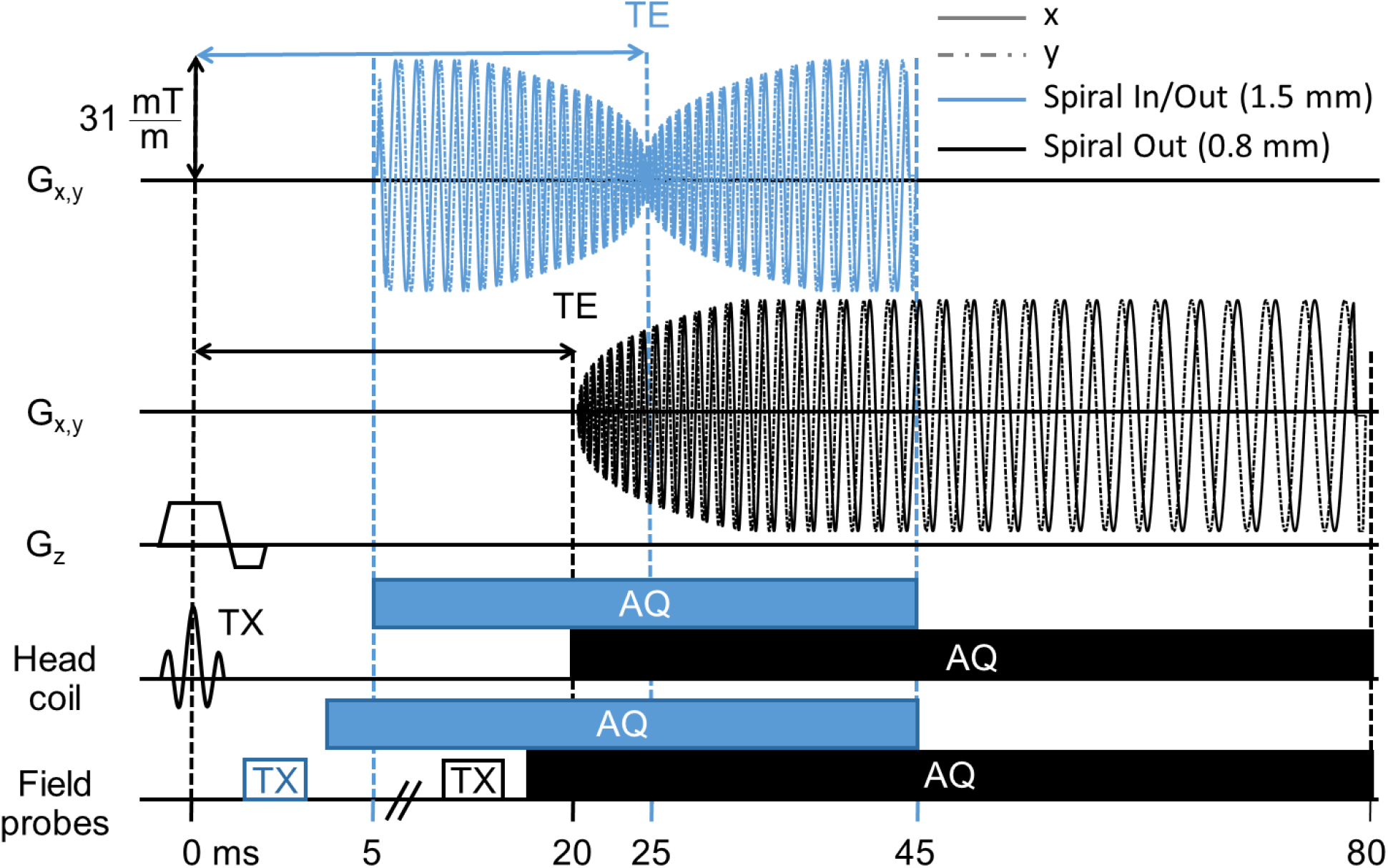
Utilized 2D single-shot spiral acquisitions (R = 4 undersampling): High-resolution single-shot spiral-out (nominal resolution 0.8 mm, black) and spiral in/out trajectory (1.5 mm resolution, blue). Depicted are the gradient waveforms (G_x_,G_y_,G_z_) as well as RF excitation (TX) and ADC sampling intervals (AQ) for both the ^1^H head coil and the ^19^F field probes used to monitor the trajectories and other concurrent encoding fields. Field probe excitation and acquisition start a few milliseconds before the spiral readout gradient waveforms.

For the fMRI experiments, visual stimulus presentation utilized VisuaStim LCD goggles (Resonance Technology Inc., Northridge, CA, USA). A vendor-specific respiratory bellows and finger pulse plethysmograph recorded subject physiology, i.e., respiratory and cardiac cycle.

### 2.2 fMRI Paradigm and Subjects

Seven healthy subjects (4 female, mean age 25.7 +/− 4.1 y) took part in this study, after written informed consent and with approval of the local ethics committee. One subject was excluded from further analysis due to reduced signal in multiple channels of the head receive array. Thus, six subjects were analyzed for this study.

The paradigm, a modified version of the one used in (Kasper et al., 2014), comprised two blocks of 15 s duration that presented flickering checkerboard wedges in complementary pairs of the visual quarter-fields. In one block, *u*pper *l*eft and *l*ower *r*ight visual field were stimulated simultaneously (condition ULLR), while the other block presented the wedges in the *u*pper *r*ight and *l*ower *l*eft quarter-fields (condition URLL). These stimulation blocks were interleaved with equally long fixation periods. To keep subjects engaged, they had to respond to slight contrast changes in the central fixation cross via button presses of the right hand. A single run of the paradigm took 5 min (5 repetitions of the ULLR-Fixation-URLL-Fixation sequence). For both of the spiral sequence designs (high-resolution spiral-out and combined spiral in/out, see next section), a single run of the paradigm was performed per subject.

### 2.3 Spiral Trajectories and Sequence Timing

Spiral fMRI was based on a slice-selective multi-slice 2D gradient echo sequence (Fig. 1) with custom-designed spiral readout gradient waveforms. For every third slice, i.e., a TR of 270 ms, concurrent field recordings were performed on the dedicated acquisition system (Dietrich et al., 2016a), with NMR field probes being excited a few milliseconds prior to readout gradient onset (Fig. 1, bottom row, (Engel et al., 2018)).

For the spiral trajectories, we selected two variants that had previously provided high-quality images in individual frames (Fig. 2 in (Engel et al., 2018)): a high-resolution case winding out of k-space center on an Archimedean spiral (spiral-out, Fig. 1, black gradient waveform), and a combined dual-image readout first spiraling into k-space center, immediately followed by a point-symmetric outward spiral (spiral in/out (Glover and Law, 2001)), Fig. 1, blue gradient waveform).

**Figure 2.**
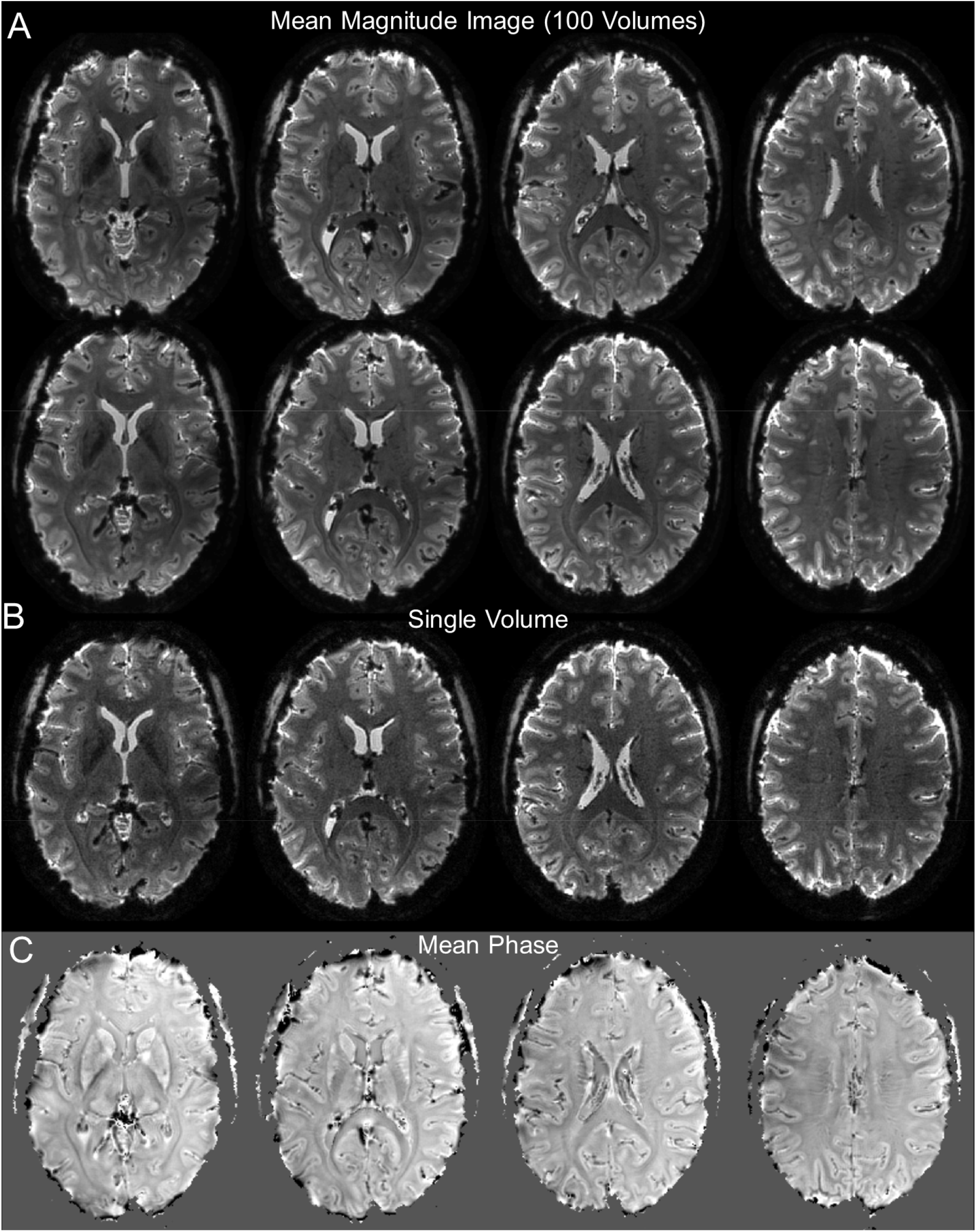
Overview of image quality for high-resolution (0.8 mm) single-shot spiral-out acquisition. (A) 8 oblique-transverse slices (of 36) depicting the time-series magnitude mean of one functional run (subject S7, 100 volumes). (B) Single-volume magnitude images for slices corresponding to lower row of (A). (C) Mean phase image over one run, without any post-processing, for slices corresponding to lower row of (A).

The spiral-out gradient waveform was designed to deliver the highest spatial resolution possible under several constraints. First, targeting maximum acquisition efficiency in 2D commands a single-shot 2D readout, because the sequence overhead, i.e., time spent without sampling k-space, accrues for each new excitation. Second, the parallel imaging capability of our receiver array at 7 T allowed for an in-plane acceleration factor of R = 4 (determining the spacing of the spiral revolutions, i.e., FOV). We based this choice on previous experience with spirals of such undersampling using this setup (Engel et al., 2018; Kasper et al., 2018), which were free of aliasing artifacts or prohibitive g-factor noise amplification. Third, the requirement of concurrent field recordings for the whole spiral readout limited its maximum duration to below 60 ms. This is the approximate lifetime of the NMR field probe signal, after which complete dephasing occurs in a subset of probes for this specific setup, governed by their T_2_* decay time of 24 ms (Engel et al., 2018) and distance from iso-center when applying higher-order shims. Finally, the gradient system specifications constrain the maximum possible resolution (or k-space excursion) of an Archimedean spiral with prescribed FOV and duration. Here, we used the optimal control algorithm by (Lustig et al., 2008) to design time-optimal spiral gradient waveforms of 31 mT/m maximum available gradient amplitude, and a 160 mT/m/ms slew rate limit, chosen for reduced peripheral nerve stimulation.

Overall, these requirements led to a spiral-out trajectory with a nominal in-plane resolution of 0.8 mm (for a FOV of 230 mm), at a total readout acquisition time (TAQ) of 57 ms. BOLD-weighting was accomplished by shifting the readout start, i.e., TE, to 20 ms.

For the spiral in/out, we followed the same design principles, targeting a minimum dead time after excitation, and a symmetric readout centered on a TE of 25 ms, slightly shorter than reported T_2_* values in cortex at 7 T (Peters et al., 2007). This resulted in a gradient waveform lasting 39 ms, with a nominal resolution of 1.5 mm for each half-shot of the trajectory.

All other parameters of both spiral sequences were shared, in order to facilitate comparison of their functional sensitivity. In particular, slice thickness (0.9 mm) and gap (0.1 mm) were selected for near-isotropic sub-mm resolution for the spiral-out case, while still covering most of visual cortex. For each slice, the imaging part of the sequence (Fig. 1) was preceded by a fat suppression module utilizing Spectral Presaturation with Inversion Recovery (SPIR, (Kaldoudi et al., 1993)).

The sequence duration totaled 90 ms per slice for the spiral-out sequence (TE 20 ms + TAQ 60 ms + SPIR 10 ms), which was maintained for the spiral in/out, even though a shorter imaging module would have been possible. To arrive at a typical volume repetition time for fMRI, we chose to acquire 36 slices (TR 3.3 s). Each functional run comprised 100 volume repetitions, amounting to a scan duration of 5.5 min.

### 2.4 Image Reconstruction

Image reconstruction rested on an expanded model of the coil signal *S_γ_* (Wilm et al., 2011), that – besides transverse magnetization *m* – incorporates coil sensitivity *C_γ_*, as well as phase accrual by both magnetostatic B_0_ field inhomogeneity (off-resonance frequency Δ*ω*_0_) (Barmet et al., 2005) and magnetic field dynamics *k_l_* expanded in different spatial basis functions *b_l_* (Barmet et al., 2008):

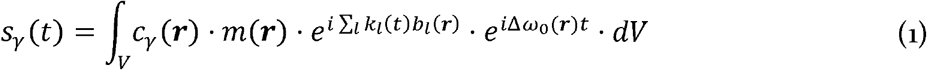

with coil index *γ*, sampling time *t*, imaging volume *v*, and location vector ***r*** = [*x y z*]^*T*^.

For 2D spiral imaging without strong higher order eddy currents (e.g., as induced by diffusion encoding gradients), this model can be computationally reduced (Engel et al., 2018) to facilitate iterative inversion. To this end, we (1) considered only field dynamics contributing to global phase *k*_0_ (such as B_0_ drifts and breathing modulation) and spatially linear phase, i.e., k-space ***k*** = [*k_x_ k_y_ k_z_*], as provided by the concurrent field recordings, and (2) restricted the integration to the excited 2D imaging plane by shifting the coordinate origin to the slice center ***r***_0_, effectively factoring slice-orthogonal field dynamics out of the integral:

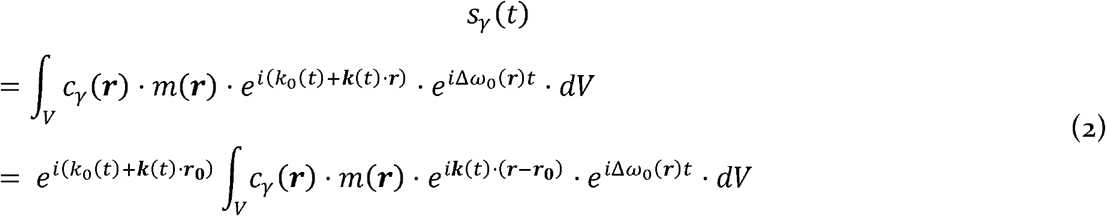

For the demodulated coil signal 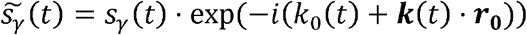, the discretized version of eq. (2) – respecting finite spatial resolution and dwell time of the acquisition system – reads as a system of linear equations

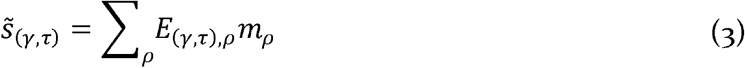

with sampling time index *τ*, voxel index 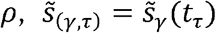, encoding matrix element 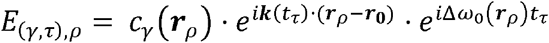, and *m_ρ_* = *m*(***r***_*ρ*_).

The matrix-vector form of eq. (3) is a general linear model,

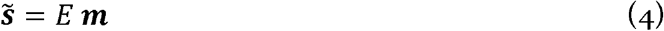

and can be efficiently solved iteratively by a conjugate gradient (CG) algorithm (Pruessmann et al., 2001; Shewchuk, 1994). As mentioned above, the restriction to first order field dynamics enables acceleration of the ensuing matrix-vector multiplications by (reverse) gridding and fast Fourier transform (FFT) (Beatty et al., 2005; Jackson et al., 1991). Off-resonance effects are also efficiently approximated by FFT using multi-frequency interpolation (Man et al., 1997).

This image reconstruction algorithm was applied equivalently to the spiral-out and spiral in/out data with a fixed number of 10 iterations and no further regularization (e.g., Tikhonov). Note, however, that for the latter both field recordings and coil data were split into their in- and out-part and reconstructed separately, yielding two images per shot.

Taken together, the in-house Matlab R2018a (The MathWorks, Natick, MA, USA) implementation of this algorithm led to total reconstruction times on a single CPU core of about 10 min per slice for the high-resolution spiral-out image and 1.5 minutes for the spiral-in or -out image. In order to reconstruct the 3600 2D images per fMRI run, reconstruction was parallelized over slices on the university’s CPU cluster. Depending on cluster load, reconstructions typically finished over night for the high-resolution spiral out, and within 2 h for the spiral in/out data.

The auxiliary input data for the expanded signal model, i.e., spatial maps for static B_0_ field inhomogeneity Δ*ω* and coil sensitivity *C_γ_*, were derived from a separate fully sampled multi-echo (ME) Cartesian gradient echo reference scan of 1 mm in-plane resolution with 6 echoes, TE_1_ = 4 ms, ΔTE = 1 ms (Kasper et al., 2018), and slice geometry equivalent to the spiral sequences. Image reconstruction proceeded as described above for this scan, albeit omitting the sensitivity and static B_0_ map terms. The latter was justified by the high bandwidth of the Cartesian spin-warp scans (1 kHz).

Sensitivity maps were then computed from the first-echo image, normalizing single coil images by the root sum of squares over all channels, while the B_0_ map was calculated by regressing the pixel-wise phase evolution over echo images. Both maps were spatially smoothed and slightly extrapolated via a variational approach (Keeling and Bammer, 2004).

### 2.5 Data Analysis

#### 2.5.1 Image Quality Assessment

The suitability of the raw imaging data for high-resolution fMRI was assessed in terms of both sensitivity and spatial specificity. No smoothing was performed for any analysis in this section.

For sensitivity, we evaluated the temporal statistics of the images, i.e., signal-to-fluctuation noise ratio (SFNR), standard deviation (SD) and coefficient of variation (CoV) maps (Welvaert and Rosseel, 2013), defined as

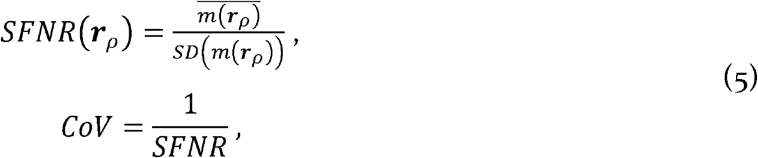

where the bar denotes averaging over volumes of a run.

Our assessment of spatial specificity was based on the ME reference scan, which exhibits a high geometric veracity due to its spin-warp nature, i.e., high bandwidth. We overlaid the contour edges (intensity isolines) of the mean (over echoes) of the ME images onto the mean spiral images 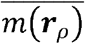 to inspect the congruency of anatomical boundaries between the scans.

To reduce the impact of subject motion on both assessments, the volumes of the fMRI time series were first realigned to each other using a 6-parameter rigid-body within-contrast registration, as implemented in SPM (Friston et al., 1996). Then, the mean ME scan was co-registered to the resulting mean realigned fMRI scan, Importantly, both operations were limited to six-parameter rigid-body registration, such that nonlinear geometric distortions between sequences were not corrected through this preprocessing step. Furthermore, to facilitate visual comparison and contour edge creation, mean ME and spiral images were bias-field corrected using unified segmentation (Ashburner and Friston, 2005).

Furthermore, for quantitative assessment, we extracted contour lines from the thresholded gray matter tissue probability maps (*ρ* ≥ 90%) retrieved by unified segmentation for both structural and functional images. To have highest contrast and resolution congruence, we compared the last echo of the 1 mm ME scan (TE 10 ms) to the 0.8 mm spiral-out scan (TE 20 ms). We computed histograms and *contour distance maps*, i.e., contours from the structural data whose color coding per voxel reflects their distance to the corresponding contour in the functional image. For the histograms, we evaluated the contour distance both over the whole imaging FOV and within an ROI of early visual cortex, the primary site of expected activation for our functional paradigm. Specifically, the ROI mask was created using the SPM Anatomy Toolbox Version 2.2b (Eickhoff et al., 2007, 2006, 2005) and combined probabilistic maps of human occipital cortex V1, V2 (Amunts et al., 2000), V3, V4 (ventral (Rottschy et al., 2007) and dorsal (Kujovic et al., 2013)), lateral occipital cortex (Malikovic et al., 2016) and V5/MT (Malikovic et al., 2007). The combined mask was warped into the individual subject geometry by the inverse deformation field retrieved through the unified segmentation mentioned above, and slightly dilated by 3 voxels to account for any inter-subject variability in visual cortex.

All computations were performed in Matlab R2019b, using the Unified NeuroImaging Quality Control Toolbox (UniQC, (Bollmann et al., 2018; Frässle et al., 2021)), and SPM12 (Wellcome Centre for Human Neuroimaging, London, UK, http://www.fil.ion.ucl.ac.uk/spm/).

#### 2.5.2 BOLD fMRI Analysis

The main goal of this analysis was to assess the functional sensitivity and spatial specificity of the spiral fMRI sequences at the single-subject level under standard paradigm and preprocessing choices. Note that, unless explicitly stated otherwise, all activation maps and their quantification (e.g., cluster extent, peak t-values) are therefore reported after smoothing.

Equivalent preprocessing steps were applied to all spiral fMRI runs using SPM12. After slice-timing correction, we employed the pipeline described in the previous section (realignment, co-registration, bias-field correction via unified segmentation). Finally, the functional images were slightly smoothed with a Gaussian kernel of 0.8mm FWHM, i.e., the voxel size of the high-resolution scan.

The general linear model (GLM) contained regressors of the two stimulation blocks (ULLR and URLL) convolved with the hemodynamic response function (HRF), as well as nuisance regressors for motion (6 rigid-body parameters) and physiological noise (18 RETROICOR (Glover et al., 2000) regressors, as specified in (Harvey et al., 2008)), extracted by the PhysIO Toolbox (Kasper et al., 2017).

To characterize functional sensitivity, we evaluated the differential t-contrasts +/− (ULLR-URLL) and report results at an individual voxel-level threshold of p < 0.001 (t > 3.22). For quantification in the results tables, we report activations under whole-brain family-wise error (FWE) correction at the cluster level (p < 0.05), given this voxel-level threshold (p < 0.001) for cluster definition. If not noted otherwise in the figure, we omitted multiple-comparison correction in the visualization of the activation maps, to study their spatial extent and specificity.

Spatial specificity of the activation was qualitatively assessed by overlaying the thresholded t-contrast maps for both contrasts onto the anatomically veridical mean ME image. We checked whether activation patterns were restricted to gray matter regions of visual cortex, as well as whether the spatial separation and symmetry of activations were linked to distinct quarter-field stimulation patterns, as expected by the retinotopic organization of visual cortex (Engel et al., 1997; Wandell et al., 2007; Warnking et al., 2002). Furthermore, we evaluated the individual contrasts for the ULLR and URLL stimulation blocks to assess the spatial overlap of their activation patterns as an alternative measure of functional specificity (since the differential contrasts cannot overlap by design).

The quantification of functional spatial specificity relied on the tissue probability maps extracted via unified segmentation from the structural scan (mean ME). To reduce the number of uncategorized voxels, we chose a liberal exceedance threshold of 60 % to define the individual tissue classes (gray matter (GM), white matter (WM), cerebrospinal fluid (CSF)). Of the remaining voxels, those that exceeded 30 % probability for two tissue classes were categorized as gray/white matter boundary (GM/WM interface) or pial surface (GM/CSF interface). All other voxels were labeled as ambiguous. We then evaluated the share of significantly activated voxels for the differential t-contrasts +/− (ULLR-URLL) after multiple comparison correction (p < 0.05 cluster-level FWE corrected with a cluster-forming voxel threshold of p < 0.001). This analysis was performed for both the whole imaging FOV and within the ROI of early visual cortex, as defined at the end of the previous section (2.5.1). We repeated this analysis for activation maps derived from unsmoothed data to study the impact of smoothing on spatial specificity.

This overall analysis procedure was performed for the spiral-out as well as the individual spiral-in and spiral-out image time series reconstructed from the spiral in/out data. As spiral in/out sequences are predominantly selected for their potential gain in functional sensitivity when combining spiral-in and spiral-out images (Glover and Law, 2001), we additionally repeated the BOLD fMRI analysis for such a surrogate dataset (“in/out combined”), but omitted the quantitative analysis of spatial specificity. We chose a signal-weighted combination per voxel (Glover and Thomason, 2004), which is considered the most practical approach for echo combination (Glover, 2012):

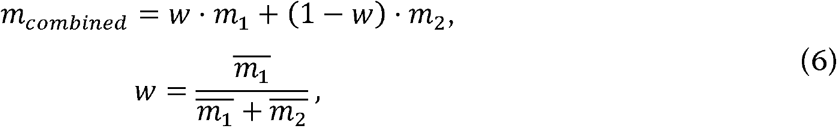

with *m*_1_ and *m*_2_ being the in-part and out-part voxel time series, respectively.

### 2.6 Code and Data Availability

Image reconstruction was performed by an in-house custom Matlab implementation of the cg-SENSE algorithm (Pruessmann et al., 2001). A demonstration of that algorithm is publicly available on GitHub (https://github.com/mrtm-zurich/rrsg-arbitrary-sense), with a static compute capsule for reproducible online re-execution on CodeOcean (Patzig et al., 2019), which were created in the context of the ISMRM reproducible research study group challenge (Maier et al., 2021; Stikov et al., 2019), albeit without the multi-frequency interpolation employed here. An example reconstruction pipeline including static B0 correction for the spiral data presented here is available on GitHub as well (https://github.com/mrikasper/julia-recon-advances-in-spiral-fmri), utilizing MRIReco.jl (Knopp and Grosser, 2021), an MRI reconstruction framework written in Julia (Bezanson et al., 2017).

Image and fMRI analyses were performed using SPM12 (https://www.fil.ion.ucl.ac.uk/spm, distributed under GPLv2) and the in-house developed UniQC Toolbox (Bollmann et al., 2018), publicly available under a GPLv3 license as a beta release within the TAPAS software collection (https://www.translationalneuromodeling.org/tapas, (Frässle et al., 2021)).

All custom analysis and data visualization scripts for this publication are available on https://github.com/mrikasper/paper-advances-in-spiral-fmri. This includes both a one-click analysis (`main.m`) to rerun all image statistics and fMRI analyses, as well as the automatic re-creation of all figure components found in this manuscript (`main_create_figures.m`), utilizing the UniQC Toolbox. More details on installation and execution of the code can be found in the `README.md` file in the main folder of the repository.

Data from this study is publicly available as part of the ETH Research Collection (doi:10.3929/ethz-b-000487412, (Kasper et al., 2021)) and described in more detail in the accompanying Data in Brief Article, according to the FAIR (Findable, Accessible, Interoperable, and Re-usable) data principles (Wilkinson et al., 2016). For one subject (`SPIFI_0007`), this includes both the reconstructed images in NIfTI format with behavioral and physiological logfiles, to validate the analysis scripts, as well as raw coil and trajectory data in ISMRMRD format (in the patient coordinate system), together with the corresponding B_0_ and sensitivity maps (NIfTI).

For the other datasets, we did not obtain explicit subject consent to share all raw data in the public domain. However, we do provide the magnetic field evolution time series as ISMRMRD files (in the scanner coordinate system) within the ETH Research Collection. Mean spiral fMRI images with corresponding activation t-maps for all subjects are also made available on NeuroVault for interactive viewing ((Gorgolewski et al., 2015), https://neurovault.org/collections/6086/).

Finally, for further data dissemination, montage views of the presented image quality metrics and statistical map overlays containing all slices and subjects are included in the supplementary materials (high-resolution spiral-out: SM 1, spiral in/out: SM 2).

## 3 Results

### 3.1 Spiral Image Quality, Congruency and Stability

In the following, we mainly present images from individual subjects (S7: Figs. 2,3, S2: Figs. 4,6,9, S3: Fig. 5). However, as illustrated by Fig. 7, as well as supplementary materials SM 1 and SM 2, results were comparable for all six analyzed datasets.

**Figure 3.**
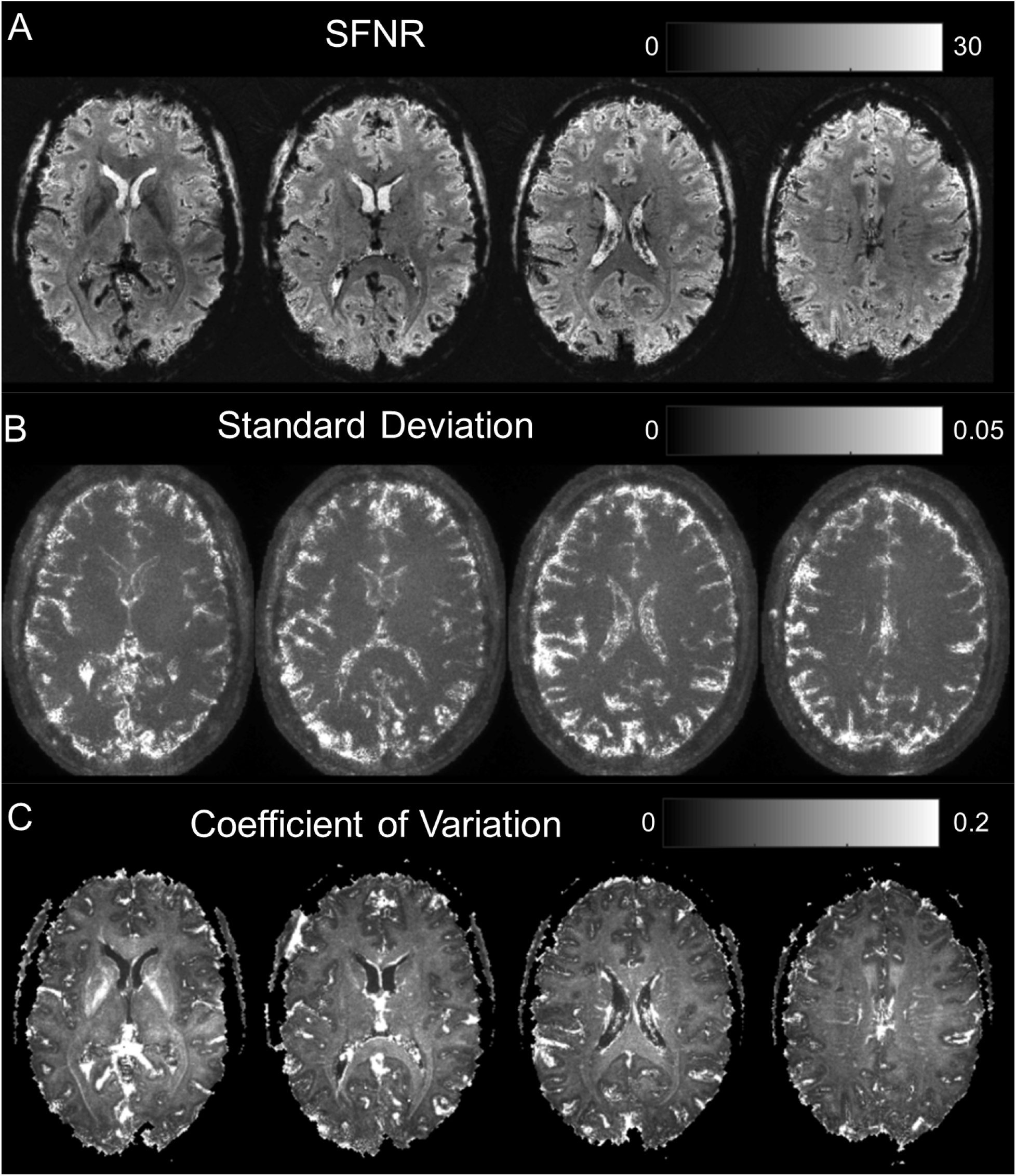
Characterization of image time series fluctuations over 1 spiral-out run (95 volumes, discarding first five). (A) Signal-to-Noise Fluctuation Ratio (SFNR) image for same slices as in Fig 2. Rather homogeneous, exhibiting sufficient SFNR levels. (B) Standard deviation (SD) image over time. Regions of high fluctuation mainly include pulsatile areas close to ventricles or major blood vessels, and cortex/CSF interfaces. (C) Coefficient of Variation (CoV) image. Inverse of (A), highlighting regions of high fluctuations relative to their respective mean. Vascularized/CSF regions appear prominently, as well as the internal capsule, due to its reduced average signal level.

**Figure 4.**
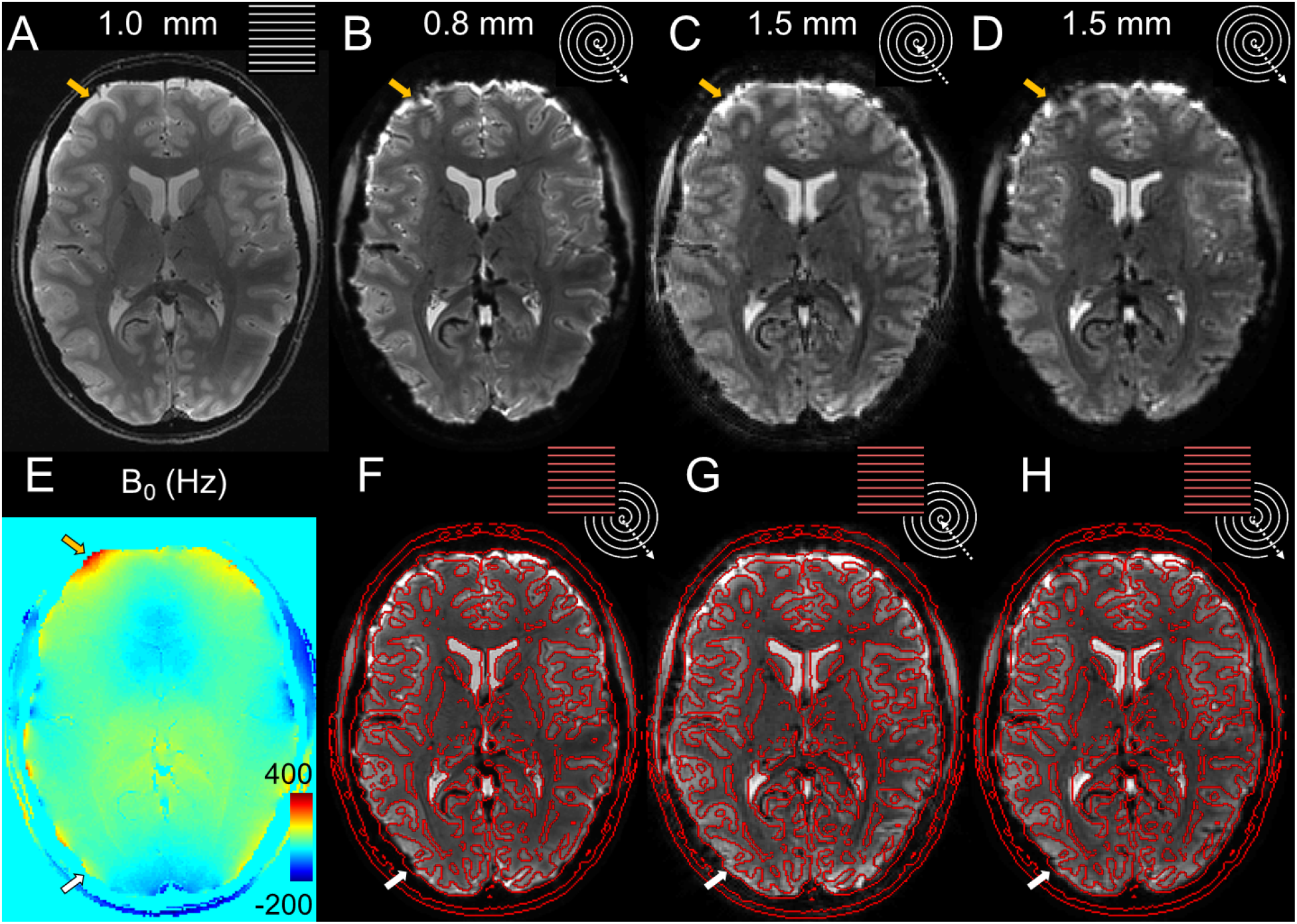
Image quality and geometric accuracy of spiral images, reconstructed with the expanded signal model. (A) Anatomical Reference: Mean multi-echo (ME) spin-warp image (1 mm resolution) (B) High-resolution (0.8 mm) spiral-out; (C) In-part of spiral in/out (1.5 mm); (D) Out-part of spiral in/out (1.5 mm). (E) B_0_-map computed from (A). (F-H) Overlay of isoline contour edges from (A) onto (B)-(D). Depicted are the mean images of a single run (subject S2, top row, B-D). The mean ME image (A), used to compute SENSE- and B_0_-map (E) for the expanded signal model, provides the anatomical reference via its contours (red lines), overlaid onto the different spiral variants (bottom row, F-H). Arrows indicate residual geometric incongruence by through-plane dephasing (white) or incomplete B_0_ mapping and correction (yellow) in the spiral-out, which are reduced in the out-part and absent in the in-part of the spiral-in/out sequence.

**Figure 5.**
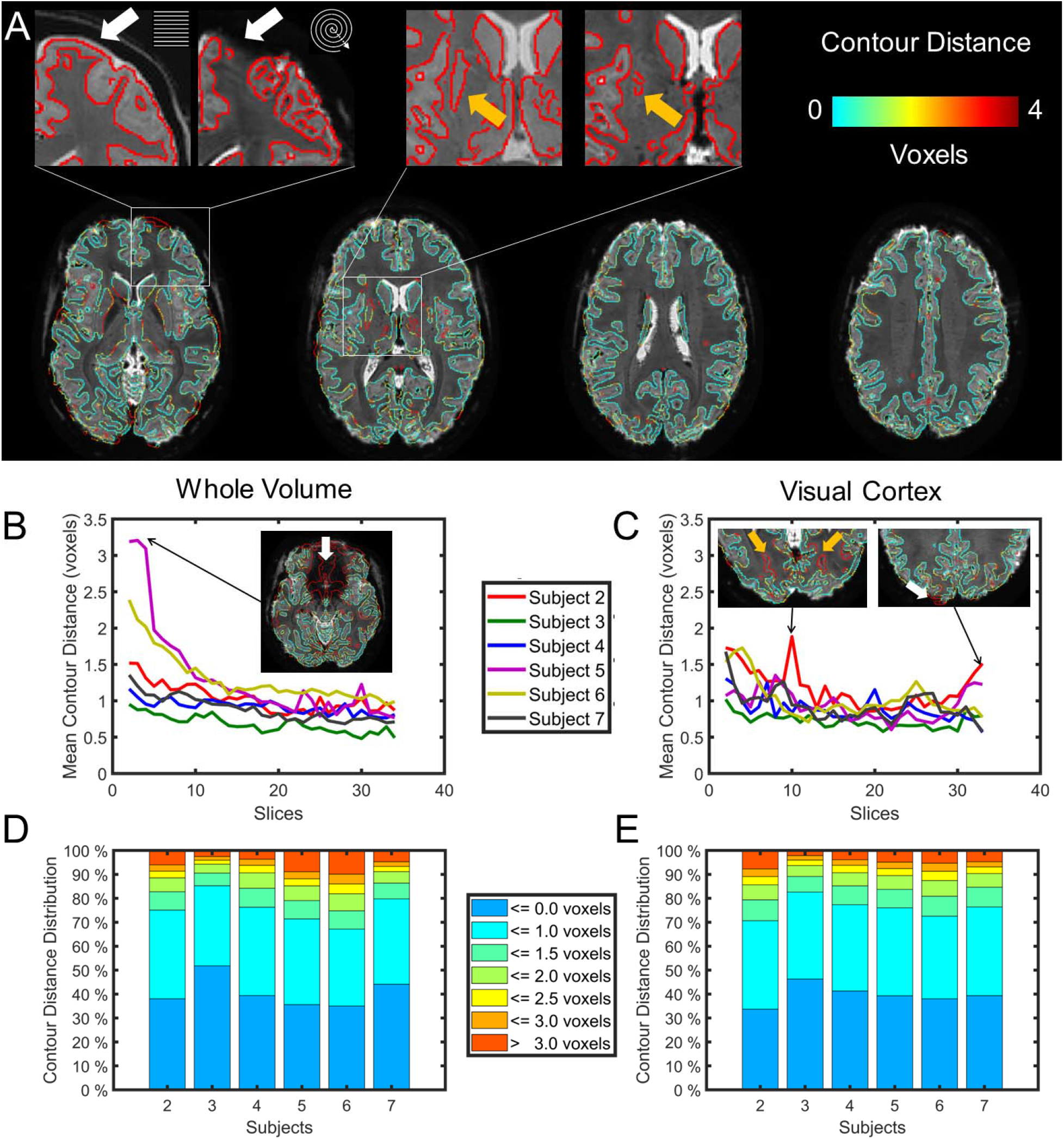
Quantification of spatial specificity in spiral images via contour distance mapping. (A) Gray matter contours extracted from tissue probability maps (threshold 90 %) of the T_2_*-weighted structural image (TE 10 ms, echo 6 of ME scan), overlaid onto mean high-resolution spiral-out image (subject S3). Color coding reflects distance to corresponding contour in segmented spiral image. Contour discrepancies are prevalent at tissue interfaces with high susceptibility gradients (left inset, white arrows), as well as areas with pronounced T_2_* contrast differences (right inset, yellow arrows). (B) Mean contour distance per slice for different subjects (averaged over all contours within each slice for the whole imaging volume). Average over subjects and slices: 1.04 ± 0.26 voxels (0.83 ± 0.21 mm). Prominent outliers (subjects S5, S6) arose in inferior slices with considerable signal loss due to through-plane dephasing (sphenoid sinus, ear canals). (C) Mean contour distance per slice for different subjects, as in (C), but restricted to contours within a mask of early visual cortex. Average over subjects and slices: 0.96 ± 0.14 voxels (0.77 ± 0.11 mm). Fewer outliers exist, mostly due to contrast differences and close to the sagittal sinus. (D) Distribution of contour distances per subject within the whole imaging volume. 41 ± 7 % of gray matter contour voxels in all subjects were strictly overlapping, with 76 ± 7 % at most 1 voxel apart and only 11 ± 5 % exceeding a distance of 2 voxels (1.6 mm). (E) Distribution of contour distances per subject, as in (D), but restricted to a mask of early visual cortex. Near-identical distribution to (D), but fewer larger outliers (≥ voxels) in some subjects (S5, S6).

**Figure 6.**
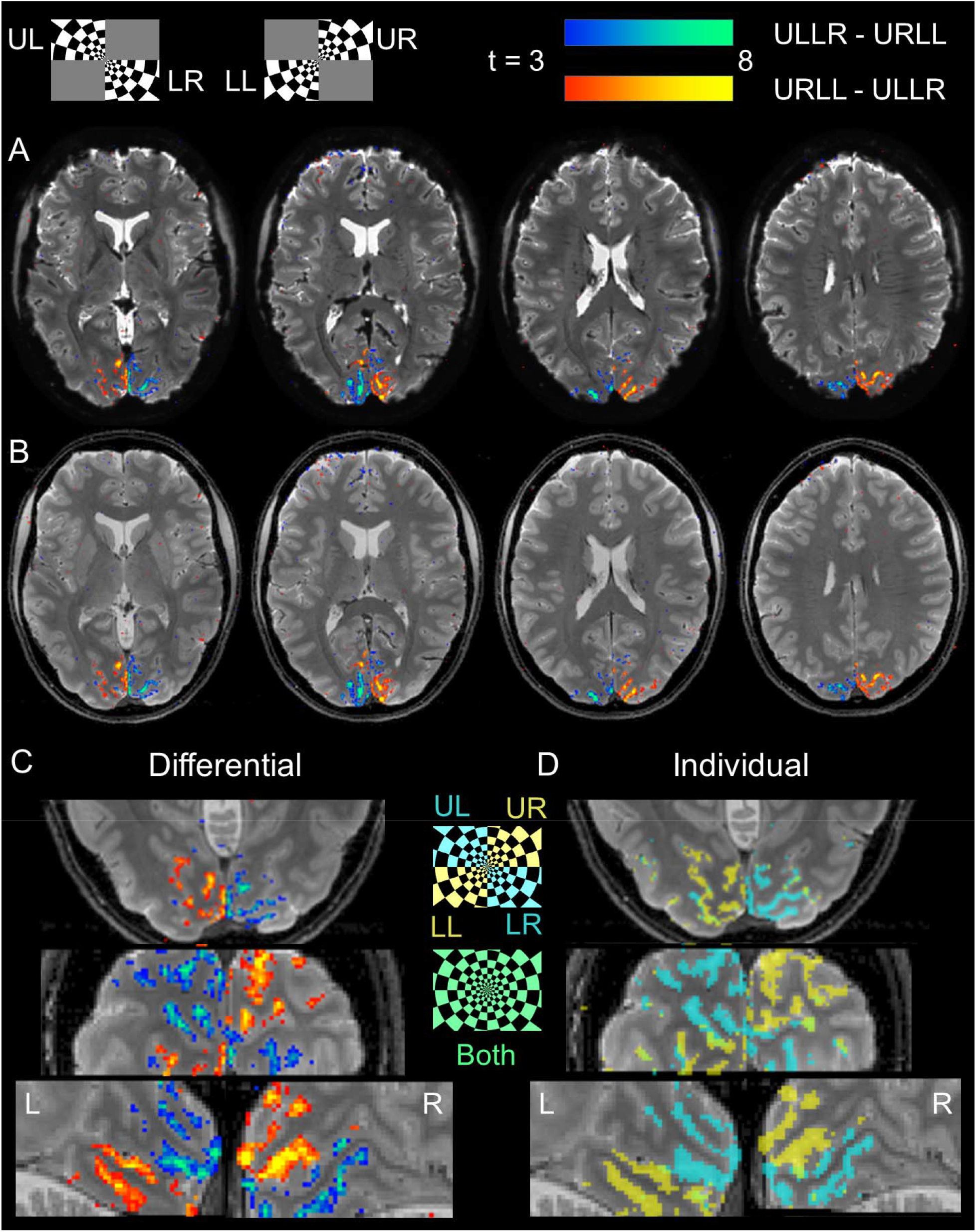
Visual Activation Maps of high-resolution (0.8 mm) spiral-out fMRI for a single subject (S2). Representative stimuli of both conditions (ULLR and URLL) are displayed at the top. (A) Overlay of differential t-contrast maps (p < 0.001 uncorrected) on transverse slices of mean spiral image (hot colormap: URLL-ULLR, cool colormap: ULLR-URLL). (B) Same contrast maps as in (A), overlaid on mean ME image as anatomical reference. (C) Zoomed-in sections of differential t-contrast maps in different orientations: transverse (top), coronal (middle) and sagittal (bottom, left (L) and right (R) hemisphere). (D) t-contrast maps for individual conditions (blue: ULLR, yellow: URLL), showing more widespread activation and high spatial specificity, i.e., little spatial overlap (green).

**Figure 7.**
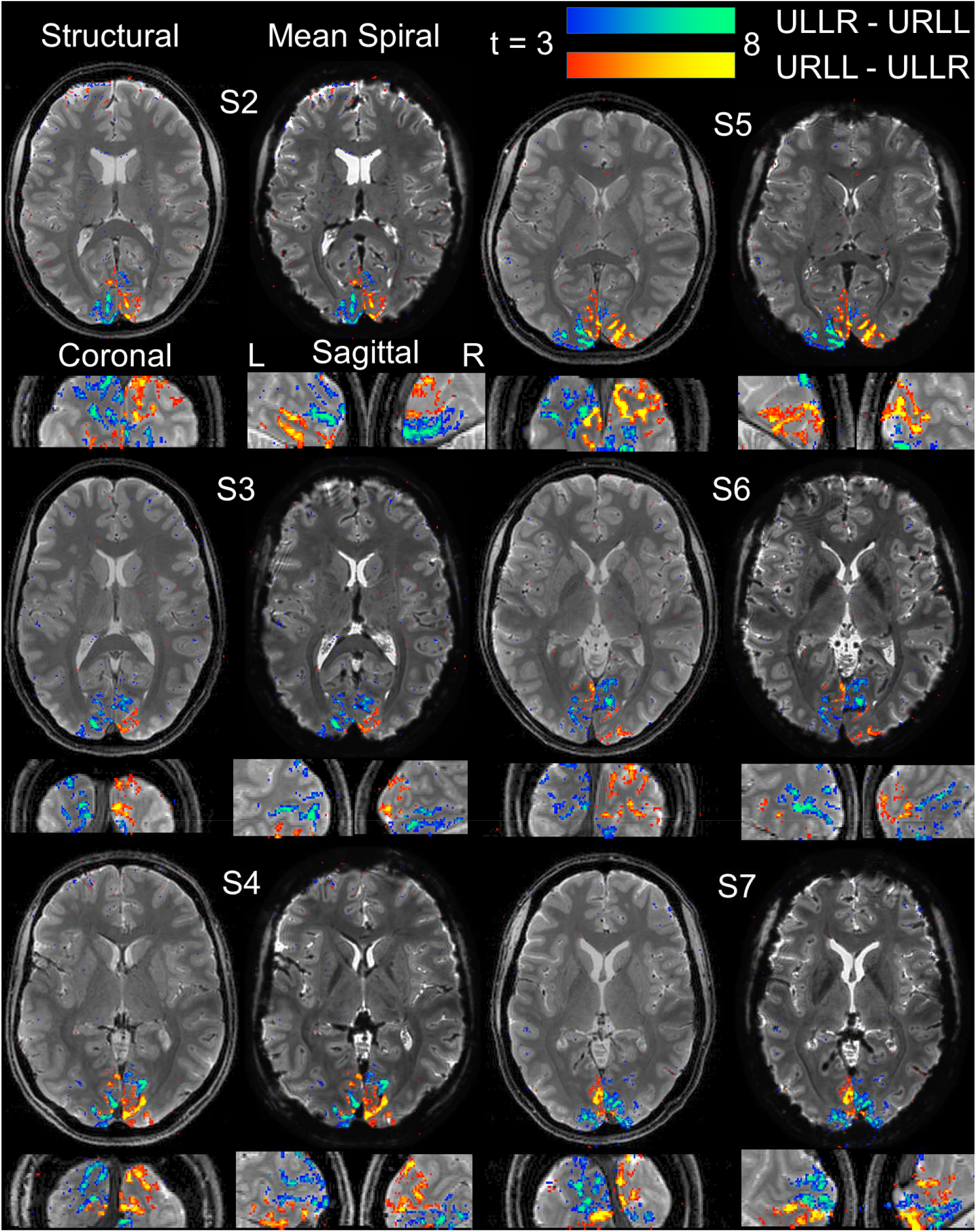
Mean spiral images and activation maps over subjects (S2-S7) for high-resolution spiral-out fMRI. For each subject, the following 4 sections are displayed, with the mean ME image as anatomical underlay: transverse, coronal and sagittal slice (for left (L) and right (R) hemisphere), each chosen for the maximum number of activated voxels (over both differential statistical t-contrasts, p < 0.001 uncorrected). To assess raw spiral data quality, the corresponding mean functional image is displayed side-by-side to the anatomical transverse slice as an alternative underlay.

**Figure 8.**
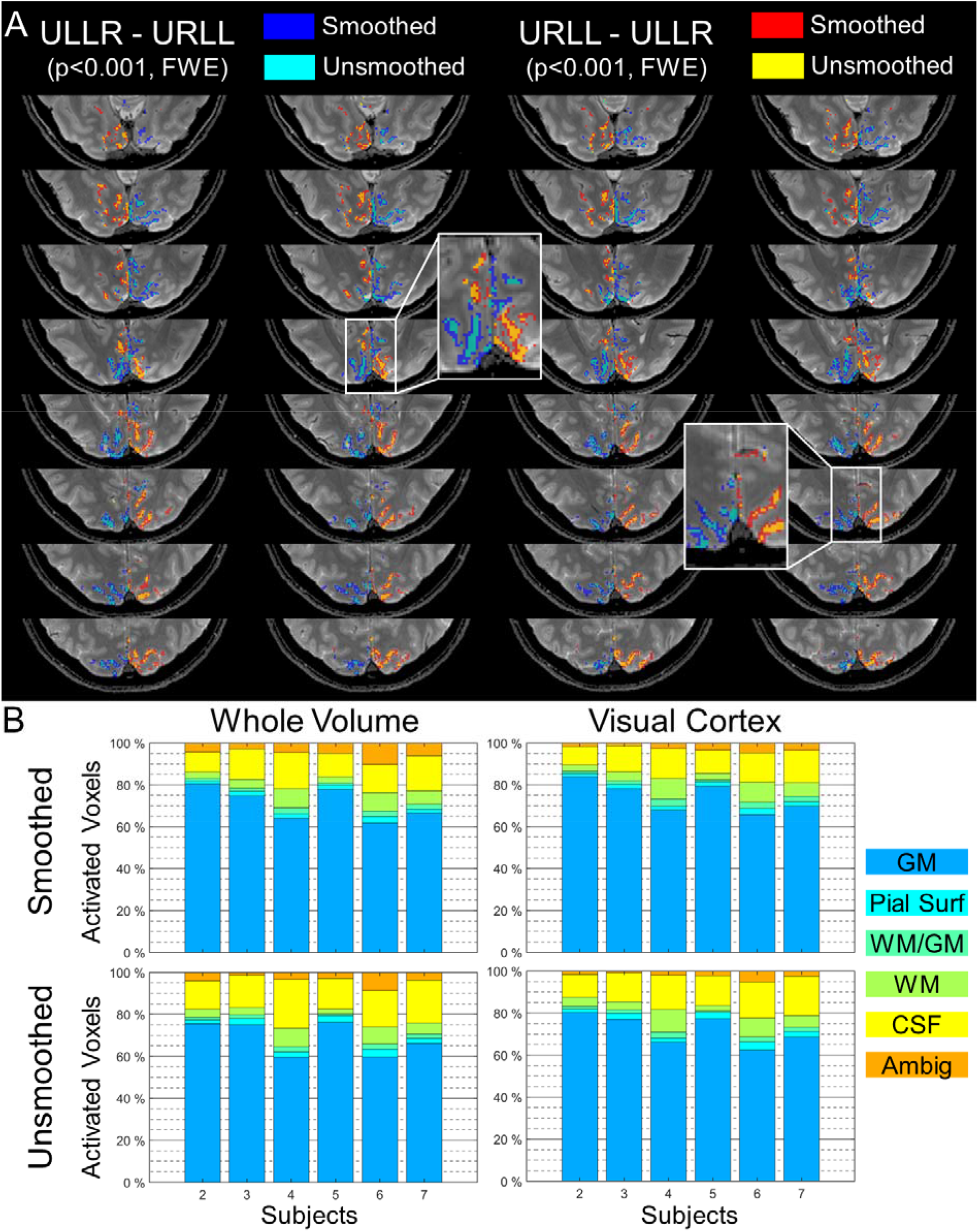
Spatial specificity of functional activation for high-resolution spiral-out fMRI. Analyses are based on significant voxels over both differential t-contrasts (+/− ULLR-URLL, p < 0.05 cluster-FWE corrected, cluster-forming threshold: p < 0.001). (A) Comparison of activation extent in smoothed (FWHM 0.8 mm) and unsmoothed data in a single subject (S2). Masks of all significant voxels are overlaid for both t-contrasts based on smoothed data (blue/red mask) as well as unsmoothed data (cyan/yellow masks). (B) Percentage of significant voxels located in relevant tissue types, analyzed for smoothed (top row) and unsmoothed (bottom row) data, as well as within whole imaging volume (left) and restricted to a mask of early visual cortex (right). Tissue types were identified by unified segmentation of the structural (mean multi-echo) image, with 60 % exceedance threshold for individual tissue classes (GM: gray matter, WM: white matter, CSF: cerebrospinal fluid) and 30 % each for interfaces (Pial surface (GM/CSF), WM/GM surface), with the remaining voxels categorized as ambiguous. On average, irrespective of smoothing and ROI, the majority (74-78 %) of activation was contained in gray matter or adjacent surfaces, with 6 % and 13-17 % residing in majority white matter and CSF voxels, respectively. Gray matter containment was highest when smoothing the data and restricting the analysis to the visual cortex, and lowest in the unsmoothed data considered within the whole imaging volume.

**Figure 9.**
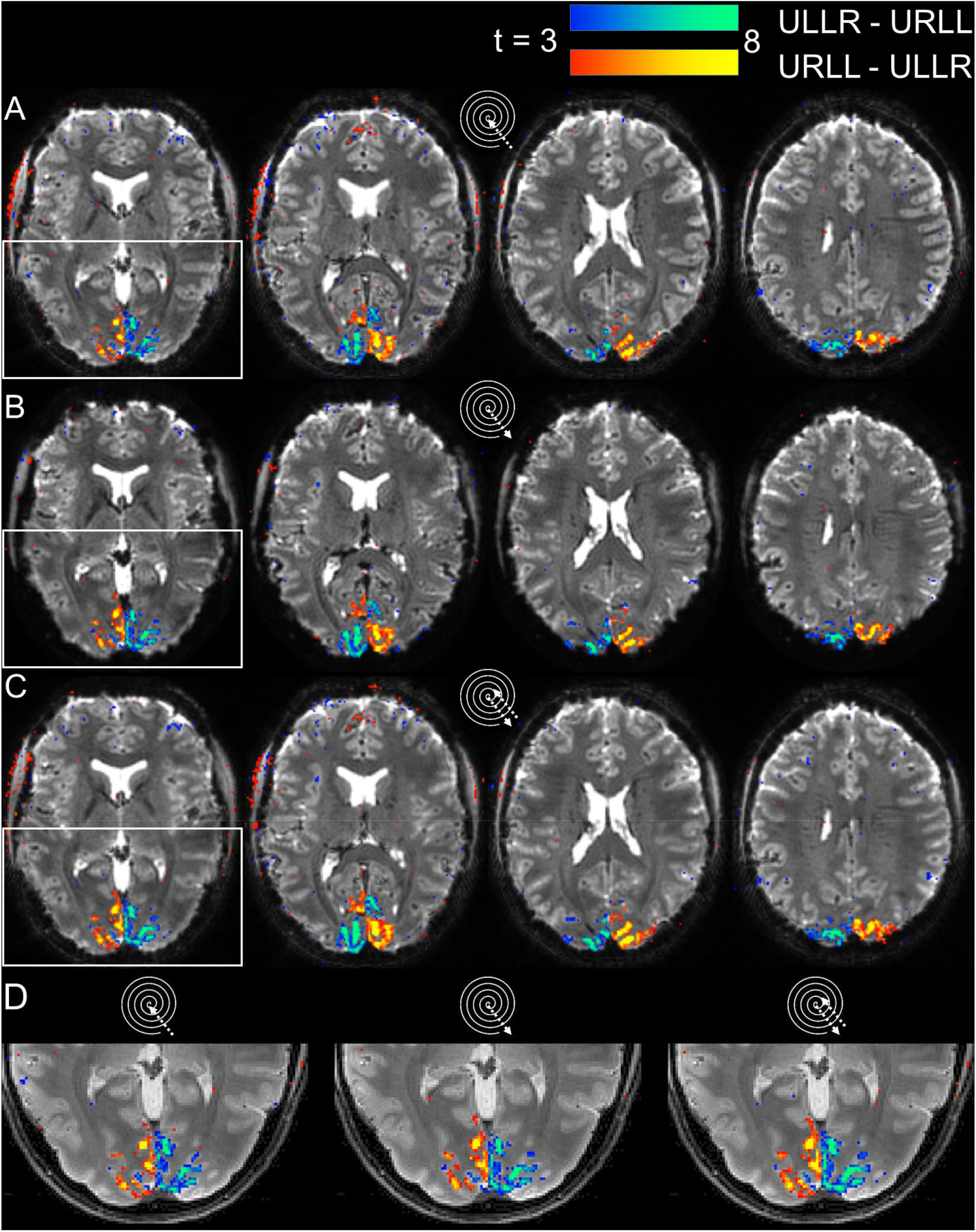
Visual Activation Maps of spiral in/out (1.5mm) fMRI run for a single subject (S2, as in Fig. 6). (A-C) Displayed are the differential t-contrast maps (p < 0.001 uncorrected) on transverse slices of the respective mean spiral images (hot colormap: URLL-ULLR, cool colormap: ULLR-URLL), based on: (A) Spiral Images reconstructed from in-part of the trajectory. (B) Spiral images reconstructed from the out-part of the trajectory. (C) Signal-weighted combination (eq. (6), (Glover and Thomason, 2004)) of images in (A) and (B). (D) Zoomed view of activation maps in leftmost slice of (A)-(C), overlaid on anatomical reference image (mean ME).

The mean images (one run of subject S7, after realignment) of the high-resolution spiral-out sequence exhibit good image quality, rich in T_2_* contrast and anatomical detail (Fig. 2A). In the center of the brain, no blurring is apparent, and anatomical boundaries can be clearly delineated, e.g, the optic radiation, down to the single-voxel extent. Moderate residual imaging artifacts (local ringing, blurring) are visible in the orbitofrontal areas, at some brain/skull boundaries, and in the vicinity of larger muscles and fat deposits, e.g., the temporal muscles. For more inferior slices, signal dropouts can be identified at typical sites of through-plane dephasing, e.g., in the temporal lobe above the ear canals (SM 1, subject 6) or in the orbitofrontal cortex (SM 1, subject 5). Individual frames of the time series exhibit similar features (Fig. 2B), though somewhat noisier, as expected because of the reduced SNR.

Interestingly, the mean of the corresponding raw phase images also contains high anatomical detail and few phase wraps (Fig. 2C), which are again located at the interface between brain and skull or close to air cavities. Note that the unwrapped appearance of the phase image is a feature of the B_0_-map based correction (Kasper et al., 2018) and does not require any postprocessing.

Mapping the temporal statistics of the spiral image time series (Fig. 3, Table 1) proves its sufficient stability for functional imaging in all slices. The SFNR images (Fig. 3A) are rather homogeneous, with mean values of 15.3 +/− 1.1 in cortical gray matter, averaged over subjects (Table 1). A slight reduction for central brain regions is visible due to the diminished net sensitivity of the receiver array. Notably, no structured noise amplification through bad conditioning of the undersampled reconstruction problem (g-factor penalty) is discernible in this area.

**Table 1.** Quantification of temporal stability and functional sensitivity of all spiral fMRI sequences. For the signal-to-fluctuation-noise ratio (SFNR, eq. (5)), the table contains mean +/− SD in a gray matter ROI over the whole imaging volume. For the t-contrast SPMs, peak t-value and number of significant voxels over both differential contrasts (+/− ULLR-URLL) are reported (p < 0.05 FWE-corrected for multiple comparisons at the cluster level with a cluster-forming threshold of p < 0.001). The last column shows relative increases to the previous sequence, i.e., the one reported in the sub-table directly above. Since resolutions differ between spiral-out (0.8 mm) and spiral in/out (1.5 mm), we compare activated volume instead of voxel count.

Advances in Spiral fMRI
| *Measure* | **Subjects** |  |  |  |  |  |  |  | **Gain vs** |  |
| --- | --- | --- | --- | --- | --- | --- | --- | --- | --- | --- |
|  | *S2* | *S3* | *S4* | *S5* | *S6* | *S7* | *Mean* | *SD* | *previous spiral* | *high-res spiral out* |
| **high-resolution spiral out** |  |  |  |  |  |  |  |  |  |  |
| SFNR_mean | 14.1 | 17.2 | 14.9 | 15.7 | 14.7 | 15.2 | 15.3 | 1.1 | - | - |
| SFNR_SD | 5.0 | 5.4 | 4.6 | 5.0 | 4.7 | 4.9 | 4.9 | 0.3 |  |  |
| SPM_T_max | 12.9 | 15.2 | 16.2 | 17.5 | 11.1 | 17.6 | 15.1 | 2.6 | - | - |
| SPM_T_nVoxels | 10957 | 6157 | 11673 | 12202 | 8682 | 12553 | 10371 | 2480 |  |  |
| SPM_T_volume (mm<sup>3</sup>) | 6832 | 3839 | 7279 | 7609 | 5414 | 7828 | 6467 | 1547 | - | - |
| **in-part spiral in/out** |  |  |  |  |  |  |  |  |  |  |
| SFNR_mean | 22.7 | 28.6 | 23.5 | 26.6 | 22.9 | 23.9 | 24.7 | 2.4 | 61% | 61% |
| SFNR_SD | 7.3 | 8.2 | 7.6 | 7.8 | 7.3 | 8.2 | 7.7 | 0.4 |  |  |
| SPM_T_max | 16.6 | 19.5 | 23.5 | 15.1 | 15.3 | 15.9 | 17.7 | 3.3 | 17% | 17% |
| SPM_T_nVoxels | 10097 | 4863 | 10221 | 6470 | 8157 | 2816 | 7104 | 2953 |  |  |
| SPM_T_volume (mm<sup>3</sup>) | 14363 | 6918 | 14540 | 9204 | 11604 | 4006 | 10106 | 4200 | 56% | 56% |
| **out-part spiral in/out** |  |  |  |  |  |  |  |  |  |  |
| SFNR_mean | 20.4 | 26.4 | 21.4 | 24.3 | 21.0 | 22.9 | 22.7 | 2.3 | -8% | 48% |
| SFNR_SD | 7.3 | 8.4 | 7.4 | 8.0 | 7.7 | 7.9 | 7.8 | 0.4 |  |  |
| SPM_T_max | 17.8 | 20.2 | 18.2 | 20.1 | 17.2 | 14.4 | 18.0 | 2.1 | 2% | 19% |
| SPM_T_nVoxels | 8976 | 6087 | 10661 | 7923 | 11606 | 3312 | 8094 | 3054 |  |  |
| SPM_T_volume (mm<sup>3</sup>) | 12769 | 8659 | 15166 | 11271 | 16510 | 4711 | 11514 | 4344 | 14% | 78% |
| **combined spiral in/out** |  |  |  |  |  |  |  |  |  |  |
| SFNR_mean | 28.3 | 35.9 | 29.5 | 33.6 | 28.7 | 30.3 | 31.1 | 3.0 | 37% | 103% |
| SFNR_SD | 9.4 | 10.9 | 10.0 | 10.4 | 9.9 | 10.7 | 10.2 | 0.5 |  |  |
| SPM_T_max | 18.4 | 19.7 | 20.3 | 17.8 | 15.8 | 18.3 | 18.4 | 1.6 | 2% | 22% |
| SPM_T_nVoxels | 11971 | 6575 | 13254 | 9331 | 11665 | 6082 | 9813 | 2985 |  |  |
| SPM_T_volume (mm<sup>3</sup>) | 17029 | 9353 | 18854 | 13274 | 16594 | 8652 | 13959 | 4247 | 21% | 116% |

The SD images (Fig. 3B) corroborate this impression, showing peak values in CSF-filled (lateral ventricles) and highly vascularized areas (insula, ACC). These noise clusters presumably stem from fluctuations through cardiac pulsation and are not specific to spiral acquisitions. However, for the raised SD values in voxels close to the cortex borders, it is unclear whether also CSF fluctuations, the BOLD effect itself, or rather time-varying blurring due to unaccounted magnetic field fluctuations contribute. This is scrutinized in the GLM analysis below. Additionally, for the CoV images (Fig. 3C), the internal capsule appears prominently, presumably due to its reduced average signal level.

In terms of spatial specificity, overlaying contour edges of the anatomical reference (mean ME spin-warp image, subject S2) (Fig. 4A) onto the mean spiral-out image suggests a geometrically very faithful depiction of the anatomical interfaces (Fig. 4B,F). Boundaries of the ventricles and gray to white matter are congruent in general, also for the visual cortex relevant to the later fMRI analyses. Some regions of the spiral-out images suffer from ringing (yellow arrow) or signal dropout (white arrow), most likely due to through-plane dephasing and incomplete correction of in-plane B_0_ inhomogeneity (Fig. 4E).

Incorporating the mean images of the spiral in/out sequence into the comparison confirms the nature of these artifacts (Fig. 4C,D,G,H). The in-part images (Fig. 4C) are devoid of these artifacts and match the anatomical reference almost completely in terms of edge contours (Fig. 4G). Only CSF/skull interfaces, for example, in frontal regions, are slightly compromised by a more global ringing, presumably from residual fat or high-intensity signal right after slice excitation, and the reversed T2* weighting in k-space for spiral-ins, amplifying high spatial frequencies of the image. The out-part of the spiral-in/out (Fig. 4D,H) constitutes a compromise between spiral-in and high-resolution spiral-out in terms of artifact-level. Its shorter readout of only 20 instead of 60 ms alleviates through-plane dephasing or incomplete B_0_ correction through inaccurate mapping.

We further quantified the spatial specificity of the high-resolution spiral-out images using contour distance mapping (Fig. 5). This measure is visualized as colored anatomical image contour on top of the mean spiral image (Fig. 5A, depicting subject S3) and shows in general good congruence of functional and structural data (0 to 1.5 voxel contour distance in most voxels). Larger deviations occurred at tissue boundaries with large susceptibility gradients, e.g., close to the frontal sinus (Fig. 5A, left inset, white arrows), as well as areas with pronounced T_2_* contrast differences between the functional and structural scan, such as subcortical gray matter (Fig. 5A, right inset, yellow arrows).

Overall, in most slices and subjects, the mean distance between corresponding contours of the structural and functional image varied between 0.5 and 1.5 voxels, with generally better congruence for more superior slices (Fig. 5B). Prominent outliers (subject 5,6, inferior slices) arose in areas with signal dropout for the spiral-out image, which was more susceptible to through-plane dephasing than the structural ME scan due to the difference in echo time (TE 2o ms vs 5-10 ms). For visual cortex in particular (Fig. 6B), the region of interest for our functional analysis, contour congruencies between 0.5 and 1 mm were most common, with fewer outliers than in the rest of the brain, driven by both contrast differences (S2, slice 10) and signal dropouts close to the sagittal sinus (S2, slice 34). Averaged over the whole volume and all subjects, the mean gray matter contour distance amounted to 1.04 ± 0.26 voxels (0.83 ± 0.21 mm), varying between 0.7 (S3) and 1.4 voxels (S5), i.e., 0.6-1.1 mm between subjects. Within visual cortex, congruence was slightly higher, with mean gray matter contour distances of 0.96 ± 0.14 voxels (0.77 ± 0.11 mm), varying between 0.7 (S3) and 1.2 (S2) voxels, i.e., 0.6-1.0 mm.

Summarizing the distribution of contour distances over all voxels in the acquisition volume (Fig. 5D), 41 ± 7 % of gray matter contours in all subjects were strictly overlapping (min (S6): 35 %, max (S3): 52 %), and 76 ± 7 % were at most 1 voxel apart (min (S6): 67 %, max (S3): 85 %), with only 11 ± 5 % of contour voxels exceeding a distance of 2 voxels (1.6 mm). This distribution was near-identical within visual cortex (Fig. 5E), with the exception of large outliers (3 voxels, i.e., 2.4 mm or more), which were reduced for individual subjects (S5, S6) from about 10 to 5 %.

### 3.2 Functional Sensitivity and Specificity

Functional sensitivity of the high-resolution spiral-out images is evident at the single-subject level (subject S2) in a differential contrast of both stimulus conditions (+/− ULLR-URLL). The corresponding t-map, overlaid on the mean functional images, reveals expected activation patterns in visual cortex (Fig. 6A). Hemispheric separation of the complementary quarter-field stimulation blocks is visible (left slice), as well as the contrast inversion from inferior to superior slices (leftmost slice vs second from left). Notably, significant activation flips between neighboring voxels occur at the cerebral fissure, suggesting spatial specificity at the voxel level.

This functional specificity is confirmed when overlaying the identical activation maps on the mean ME image as anatomical reference (Fig. 6B), again demonstrating the good alignment of functional and structural data seen in the previous subsection (Figs. 4,5). Clustered activation is almost exclusively constrained to gray matter with no extension into adjacent tissue or skull. Note that no multiple comparison correction was performed for visualization, in order to be more sensitive to such effects, at the expense of occasional false-positive voxels throughout other brain areas.

Gray-matter containment and retinotopic organization of the activation can be further corroborated in the zoomed-in sections of visual cortex for transverse, coronal and sagittal orientation (Fig. 6C). Additionally, we evaluated the ULLR and URLL blocks individually (Fig. 6D), because differential contrasts, by design, do not allow for spatial overlap between significant activation of both conditions. In the individual contrasts, the identified portion of activated visual cortex appears larger, but is still very well restricted to cortical gray matter. Few overlaps exist, and, again, contrast switches between adjacent voxels, pointing to spatial specificity at the prescribed sub-mm resolution.

These findings are reproducible over subjects (Fig. 7). Importantly, similar image quality and geometric congruency are accomplished in all subjects. To verify, we show both the mean spiral and the anatomical ME reference image of the corresponding transverse slice as underlays for the differential activation patterns. Some subjects exhibit more frontal blurring artifacts and dropouts (S5, S6, S7) due to different geometry of the air cavities. Still, the retinotopic organization of visual cortex is recovered in all subjects, as visualized in the zoomed coronal and sagittal views. Existing differences of the specific activation patterns are within the expected range of variability in subject anatomy and task engagement. Quantitatively, peak t-values reach 15.1 on average for the differential contrasts, with a standard deviation of 2.6, i.e. 17 %, over subjects (Table 1). Activation clusters comprise 10371 +/− 2480 voxels (after FWE-multiple comparison correction at the cluster level, p < 0.05), i.e., 6467 +/− 1547 mm^3^.

Because traditional Gaussian smoothing is frequently omitted in ultra-high resolution fMRI studies (e.g., for laminar fMRI), we assessed its impact on our results. We repeated the statistical analysis for all subjects to create a version of Figure 7 based on unsmoothed data (supplementary material SM 3). For one particular subject (S2), we also juxtaposed spatial characteristics of the statistical t-maps in an animated slide show by varying significance thresholds for smoothed and unsmoothed data, as well as cropping the spiral k-space data to 1 mm resolution before reconstruction (supplementary material SM 4). Overall, spatial smoothing increased overall CNR (higher t-values) and extent of activation clusters that were already discernible in the unsmoothed data. The activation clusters of the smoothed data resemble those of unsmoothed data at lower thresholds, but with fewer single-voxel activation sites. When overlaying activation masks of both differential t-contrasts (+/− ULLR-URLL) after cluster-level multiple comparison correction (p < 0.05 cluster-FWE, cluster-forming threshold: p < 0.001) for this subject directly, we observed two distinct effects of the employed moderate smoothing (FWHM 0.8 mm) (Fig. 8A): on the one hand, cluster extent may increase isotropically by about one voxel (left inset), consistent with a loss in spatial specificity. On the other hand, clusters can expand by several voxels along the cortical ribbon after smoothing (right inset), suggesting that increased sensitivity by averaging of thermal noise can lead to functionally more plausible activation patterns.

To quantify functional specificity, we evaluated the tissue type of all significantly activated voxels, and assessed the impact of smoothing on this measure (Fig. 8B), with tissue type based on the unified segmentation results of the structural data (mean ME). For smoothed data and considering the whole imaging volume (top left), 71 ± 8 % of all significantly activated voxels resided in GM (mean and standard deviation over subjects and whole volume), 2.2 ± 0.5 % and 1.9 ± 0.9 % at pial surface and GM/WM interface, respectively, 5.7 ± 2.7 % in WM and 13 ± 3 % in CSF, while for the remaining 5.5 ± 2.5 % of significant voxels, tissue type could not be determined unambiguously. Gray matter containment dropped by about 2 % in the unsmoothed data (bottom left), presumably due to randomly distributed false positives. This small difference is preserved when restricting the analysis to the ROI of early visual cortex (right column), in which gray matter containment is about 3 % higher for both smoothed and unsmoothed data. This indicates that the impact of smoothing on this quantification of functional specificity was small on average, and that the activation containment was comparable in early visual cortex and the whole imaging volume.

### 3.3 Spiral In/Out Analysis and Echo Combination

We continue to present data from the same subject (S2) as in the high-resolution case, to facilitate comparison. All findings are generalizable over subjects, and we provide mean, SD, SFNR and t-maps of all slices for further dissemination in the supplementary material (SM 2).

Overall, the differential t-contrast maps for the spiral in/out data resemble the activation patterns of the high-resolution spiral-out case (Fig. 9). This holds for all three derived in/out time series, i.e., the separate reconstructions of the in-part and the out-part, as well as their combination in the image domain via signal-weighted averaging (“in/out combined”).

In terms of functional sensitivity, the in/out sequence provides higher peak t-values and cluster extents in the differential t-contrasts compared to the high-resolution spiral-out, as expected due to the larger voxel size and consequential higher SFNR (Table 1). For example, the in-part itself provides a 61 % SFNR increase in gray matter (averaged over subjects), 17 % increased maximum peak t-value, and 56 % increase in significantly activated gray matter volume (Table 1, rightmost column).

Comparing the out- to the in-part of the spirals, SFNR is slightly decreased in the out-part (8 %), while the situation is reversed for the t-maps, with 2 % increase in peak t-value and 14 % increase in cluster extent, compared to the spiral-in. This suggests that higher T_2_*-sensitivity of the spiral-out causes both effects, by both amplifying signal dropouts and BOLD signal.

The signal-weighted echo combination (eq. (6), (Glover and Thomason, 2004)) provides the highest functional sensitivity of the three in/out time-series, having a 25 % increased SFNR compared to the in-part, and 37 % increase compared to the out-part. This translates into an average increase in peak t-value of 2 % and significant cluster extent of 21 %, compared to the out-part alone. This is in line with previous findings for high-resolution multi-shot spiral data (Singh et al., 2018) at 3 T, which also reported contrast-to-noise ratio (CNR) increases for signal-weighted spiral in/out combinations on the order of 25 %. However, it falls somewhat short of the 54 % increase in CNR reported originally for low-resolution single-shot spiral in/out combination (Glover and Thomason, 2004, p. 866).

In terms of spatial specificity, all activation patterns exhibit a good congruency to the anatomical reference, as evident from a close-up overlaid onto the mean ME image (Fig. 9D). In general, this visualization confirms the overall impression that the echo combination increases CNR throughout visual cortex, rather than just in regions of higher dephasing. Remarkably, there seem to be more false positive clusters for the spiral-in than in all other spiral variants (Fig. 9A), in particular close to the temporal muscle, presumably due to the ringing mentioned above.

## 4 Discussion

### 4.1 Summary

In this work, we demonstrated that recent advances in high-resolution, single-shot spiral imaging (Engel et al., 2018) can be deployed to fMRI. The typical drawbacks of spiral fMRI, which have so far limited its routine use, were overcome by an expanded signal model, accurate measurement of its components, and corresponding iterative image reconstruction (Barmet et al., 2005; Pruessmann et al., 2001; Wilm et al., 2011).

Specifically, time series of high image quality and stability were obtained that exhibited geometric congruency to anatomical scans without the need for post-hoc distortion correction. Notably, also the corresponding phase images exhibit high raw data quality (without any postprocessing, e.g., phase unwrapping), and suggest the suitability of spiral acquisition for novel phase- or complex-value based fMRI analysis workflows (Balla et al., 2014; Bianciardi et al., 2014; Calhoun et al., 2002; Menon, 2002).

The functional sensitivity of spiral readouts was confirmed by observing typical activation patterns in response to an established visual quarter-field stimulation. While a consensus on how to assess spatial specificity for fMRI is lacking, several indicators point to a localization capability in the sub-mm range for our data. First, the distance of gray matter contours in spiral and structural MRI data were at most one voxel (0.8 mm) apart in the vast majority of voxels per subject (76 %). Second, the activation patterns of different stimulus conditions could be discriminated in neighboring voxels of 0.8 mm nominal resolution (Fig. 6). Third, the vast majority (75 %) of significant activation sites were contained within gray matter or at its boundaries, suggesting a limited impact of artifactual blurring.

Finally, we demonstrated the versatility of this approach to spiral fMRI with a combined in/out readout at a more typical resolution (1.5 mm). Here, the high acquisition efficiency of the spiral allowed to measure two images per shot, increasing CNR by about 20 %. The observed discrepancy to previously reported gains of more than 50 % for signal-weighted echo combination (Glover and Thomason, 2004) is in line with recent spiral fMRI studies (Singh et al., 2018). It might result from higher target resolution and static off-resonance correction employed in our study and (Singh et al., 2018), compared to the original work. The increased image congruency and smaller dephasing effects between the in- and out-part compared to low-resolution spirals may reduce the impact of echo combination. Still, more sophisticated combination of echo images (Glover and Thomason, 2004), or of k-space data during reconstruction (Jung et al., 2013) could result in further SNR increases.

In summary, the presented advances render spiral fMRI an attractive sampling scheme that delivers on the long-time postulate of high acquisition efficiency without compromising image quality. Here, the spatiotemporal application domain of fMRI on a standard gradient system was enhanced by acquiring a 230×230×36 mm FOV brain image at 0.8 mm nominal in-plane resolution (i.e., a matrix size of 288×288×36) while maintaining a TR typical for high-resolution fMRI (3.3 s). This corresponds to an acquisition efficiency of about 900,000 resolved voxels per second.

To our knowledge, this is the highest acquisition efficiency reported for 2D spiral fMRI to date (see Table SM 5 for a comparison of sequence parameters in several spiral fMRI studies), as well as the first high-resolution spiral fMRI study at ultra-high field. In combination with the presented evidence for geometric accuracy, this makes the presented spiral-out sequence an attractive candidate for high-resolution applications of fMRI, studying the functional sub-organization of cortex, e.g., in laminae or columns (Cheng et al., 2001; Feinberg et al., 2018; Fracasso et al., 2016; Huber et al., 2017a; Kok et al., 2016; Koopmans et al., 2010; Lawrence et al., 2018; Martino et al., 2015; Muckli et al., 2015; Siero et al., 2011; Uğurbil et al., 2013; Yacoub et al., 2008).

### 4.2 Effective Resolution, Spatial Specificity and Congruency

The claim of high acquisition efficiency for fMRI hinges on whether the acquired voxels effectively resolve distinct activation. This question of *effective* functional resolution arises for any fMRI protocol and comprises global aspects, such as PSF broadening, as well as more localized effects concerning geometric congruency and spatial specificity of the activation mapping.

The circular k-space coverage of spirals leads to a broadening of the PSF main lobe by 17 % compared to Cartesian k-space coverage in EPI (1.4 vs 1.2 times the voxel size (Qin, 2012)). Furthermore, any sequence with long readout duration encounters considerable T_2_* signal decay along the trajectory, which manifests as a filter in image domain. For spiral-in images, this emphasizes higher spatial frequencies, while the effect on spiral-out images is reversed, leading to blurring. Based on typical brain tissue relaxation times at 7T, we adapted previous simulations of this effect for a similar high-resolution spiral-out protocol (Fig. 7 in Engel et al., 2018). There are diminishing returns for investing more acquisition time to achieve higher in-plane resolution, but a net gain remains at our chosen readout duration of 60 ms. Effectively, the FWHM of the PSF due to T_2_* blurring corresponds to a voxel size smaller than 1 mm for the targeted 0.8 mm nominal resolution, while at 40 ms readout duration actual voxels are larger than 1.1 mm for a targeted 1 mm nominal resolution. Still, choosing shorter readouts with slightly coarser resolution in favor of sampling more slices within the given TR might deliver overall higher acquisition efficiency in this case. Finally, static B_0_ inhomogeneity also manifests as spatially varying blurring or ringing in spiral imaging, because off-resonance induces broadening of the PSF main lobe, as well as amplification of its side lobes (Bernstein et al., 2004, Chap. 17; Fig. 6 in Engel et al., 2018; Man et al., 1997). As long as B_0_ inhomogeneity is properly mapped and included in the signal model, this effect is mitigated by the iterative image reconstruction utilized in this work.

In our experimental data, we found that the spatial specificity of spiral fMRI is very high in about 75-80 % of the voxels, as indicated by both contour distance mapping (Fig. 5) and gray matter containment of activation (Fig. 8). While these quantifications provide rough estimates of functional spatial specificity in fMRI, there is no consensus on such quantification in the community, and the absolute values reported here are hard to compare to previous work. We hope that through our sharing of the code, this methodology may provide future reference points. Furthermore, the utilized measures are in themselves imperfect assessments of spatial specificity, and might underestimate the achieved spatial specificity in our data. First of all, both analyses relied on unified segmentation (Ashburner and Friston, 2005) to extract tissue probability maps, which in principle should work contrast-independently. However, the employed default parameter settings may not be optimal to model bias fields at 7 T or voxel intensity distributions of multi-echo GRE scans with only partial brain coverage. Additionally, because the contrast in our functional and structural scans were not equivalent (e.g., TE 20 vs 10 ms), contour distance mapping reflects, to a certain extent, differences in contrast rather than geometry.

Finally, relying on a perfect retinotopic organization of visual cortex for assessing functional spatial specificity has also limitations. For example, receptive fields can cross the vertical meridian (especially in higher visual areas), such that differential contrasts of quarterfield stimulation may not flip between adjacent voxels (Fig. 6C). Similarly, overlapping voxel activation or single-voxel “false positives” (Fig. 6D) may indicate imperfections in the visual field leading to non-compact retinotopic representations, rather than losses in spatial specificity.

Another contentious point concerning spatial specificity is our choice of moderately smoothing the data (FWHM 0.8 mm) before statistical parametric mapping. While smoothing is a standard preprocessing step in the majority of fMRI studies, high-resolution applications, such as retinotopic mapping or layered fMRI analyses, frequently abstain from it or use more spatially informed averaging *methods* (Blazejewska et al., 2019). From a conceptual point of view, smoothing alters the target PSF of the imaging process in multiple ways that affect the effective resolution: On the one hand, a Gaussian filter, as employed here, broadens the PSF main lobe (PSF) reducing spatial specificity. On the other hand, it suppresses PSF side lobes and thus contamination by remote locations, which enhances spatial specificity. The opposite is true for an edge or high-pass filter, as implemented, for example, by multiplying the inverse of the T_2_* decay curve onto the k-space data. Overall, resolution can be re-negotiated by appropriate filtering, which has to be adapted to the specific application, in order to provide optimal specificity.

For our spiral data in particular, the decision to smooth was governed by another goal of filtering, namely, recovering sensitivity. Theoretically, sensitivity is maximized by a matched filter resembling the spatial activation extent, which is traditionally assumed to be a Gaussian for fMRI (Friston, 2007, Chap. 2; Kasper et al., 2014), but has, to our knowledge, not been determined for the ultra-high-resolution regime in question here. Our choice of smoothing with an FWHM equivalent to the voxel size (instead of 2-3 times the voxel size as in standard fMRI) therefore constitutes a compromise between sensitivity and specificity, motivated by the noise levels in our raw data, the short run duration (5.5 min) and by having only one run acquired per subject and spiral sequence. This hampered our ability to quantify whether the investment into longer readouts of nominal 0.8 mm resolution, compared to 1 mm, translated into more spatially specific activation (supplementary material SM 4). More temporal averaging via longer and more numerous functional runs would allow to address this important research question in future studies.

### 4.3 General Applicability and Limitations of Spiral Imaging Advances

#### 4.3.1 Rationale

Increasing acquisition efficiency for high-resolution single-shot spirals while maintaining depiction quality, as presented here, resulted from the favorable interplay of the expanded signal model components: the encoding field dynamics with long-readout spirals, static B_0_ inhomogeneity characterization, and parallel imaging with iterative reconstruction enabling undersampling.

For deploying this advanced spiral functional imaging to other sites and systems, it is important to evaluate how individual aspects of the approach contribute to its overall performance, and to assess the generalizability of our findings. This includes both the impact of model and system components, as well as the utilized methodology for their characterization, in relation to possible alternatives and extensions.

#### 4.3.2 Magnetic Field Monitoring

In terms of availability, the concurrent field monitoring hardware employed in our approach (Barmet et al., 2008; Engel et al., 2018; Kasper et al., 2018; Wilm et al., 2017, 2011), is probably the scarcest resource across sites. It serves to characterize both the reproducible and irreproducible imperfections of the encoding magnetic fields.

For reproducible field effects, such as the actual spiral trajectory performed by the system and its induced eddy currents, previous work has shown that their characterization is often required to avoid severe image artifacts (Engel et al., 2018, Fig. 6; Vannesjo et al., 2016). This, however, might vary between systems, as successful spiral image reconstructions based on nominal trajectories have been reported (Kurban et al., 2019; Singh et al., 2018). If image artifacts arise from reproducible trajectory imperfections, they could be measured without concurrent field monitoring hardware by calibration approaches in a separate scan session (Bhavsar et al., 2014; Duyn et al., 1998; Robison et al., 2019, 2010). For more flexibility, the gradient impulse response function (GIRF) to arbitrary input trajectories can be modelled from such data under linear-time invariant system assumptions (Addy et al., 2012; Campbell-Washburn et al., 2016; Rahmer et al., 2019; Vannesjo et al., 2014, 2013). The required field measurements for these calibrations may either rely on dedicated NMR-probe based field cameras (Barmet et al., 2008; Vannesjo et al., 2014, 2013; Zanche et al., 2008) or on off-the-shelf NMR phantoms (Addy et al., 2012; Duyn et al., 1998; Rahmer et al., 2019), with certain trade-offs to measurement precision and acquisition duration (Graedel et al., 2017).

For the dynamic field effects, induced by the system (e.g., drifts through gradient heating), as well as the subject (e.g., fluctuations with the breathing cycle or limb motion), few studies have analyzed their impact on spiral fMRI time series (Pfeuffer et al., 2002). In principle, the concurrent field monitoring and reconstruction employed here incorporated changes of global off-resonance and k-space with the bandwidth of the trajectory measurement of about 4 Hz (monitoring every third slice). As we did not observe any conspicuous problems in the time series statistics, for example, SFNR drops, nor any indication of time-dependent blurring, which would be the spiral equivalent to apparent motion in phase encoding direction observed in EPI (Bollmann et al., 2017; Power et al., 2019), this approach presumably addressed the majority of field fluctuations present in our data. This is in line with previous results of breathing-induced field fluctuations reported at 7T in spiral fMRI of only a few Hz for healthy subjects and normal breathing (Pfeuffer et al., 2002). An in-depth analysis of these effects is beyond the scope of this paper, as it would, for example, entail quantitative comparisons with nominal or GIRF reconstructions (Vannesjo et al., 2016), as well as simulating the impact of different measured field components on image time series, similar to work previously conducted for EPI (Bollmann et al., 2017; Kasper et al., 2015). As we do believe that this investigation is relevant to the neuroimaging community, we provide the field dynamics of all spiral-out fMRI runs in ISMRMRD format for further scrutiny.

Note, however, that the dataset in itself might not be representative for assessing the utility of concurrent field monitoring for spiral fMRI. In terms of system fluctuations, we did have a challenging gradient duty-cycle (with gradients switched on during 70 % of the sequence, 50 % of the time at amplitude maximum), leading to substantial heating of 15 degrees throughout the 5.5 min high-resolution spiral-out sequence, as measured using 5 optical sensors cast into the gradient coil (Dietrich et al., 2016b). This actually limited the duration of our functional runs. While system fluctuations might thus be particularly pronounced in our data, the subject-induced fluctuations will be moderate, because all volunteers were young, healthy individuals instructed to lie still throughout the session. This is reflected in the small mean framewise displacement (FD) encountered in all subjects (mean FD and standard deviation over subjects 0.09 ± 0.04 mm, see SM 6 for motion and FD traces). For a comprehensive assessment, instructed limb motion and deep breathing, a range of BMIs and body shapes would have to be included in the design of the study, similar to evaluations of field effects on structural T_2_* imaging (Duerst et al., 2016). Finally, in terms of the chosen imaging FOV covering the visual cortex, dynamic field effects will be at an intermediate level, with maximum fluctuations expected in inferior regions closer to the chest (brainstem, cerebellum) and minimum effects near the top of the head (e.g., motor cortex).

If dynamic field effects constitute a significant artifact and noise source for spiral fMRI time series, in lieu of field monitoring, alternative correction methods comprise dynamic off-resonance updates or higher-order field navigators (Pfeuffer et al., 2002; Splitthoff and Zaitsev, 2009), as well as gradient response models that incorporate time-courses of gradient coil temperature (Dietrich et al., 2016b; Stich et al., 2020) or current measurements (Nussbaum et al., 2019; Rahmer et al., 2021).

#### 4.3.3 Static B_0_ Inhomogeneity

To characterize static B_0_ inhomogeneity, we acquired a Cartesian multi-echo gradient echo scan with rather high resolution (1 mm). Including this information in the signal model has previously been crucial to maintain spatial specificity in spiral imaging at 7T (Engel et al., 2018, Fig. 6; Kasper et al., 2018, Fig. 7).

The B_0_ maps obtained in this work exhibited considerable inhomogeneity, even after 3^rd^ order shimming of the targeted 4 cm oblique-transverse slab of the brain including the visual cortex. For the B_0_ map of a single subject (SPIFI_0007) provided in the accompanying Data In Brief article (see “Code and data availability” section), 20 % of brain voxels were more than 50 Hz off-resonant, which – if uncorrected for – would incur blurring with FWHMs of several voxels.

Thus, some form of static B_0_ inhomogeneity correction seems indispensable for providing high spiral image quality, and to correct this at the reconstruction stage via inclusion into the expanded signal model proved sufficient for most of the imaged brain slices. However, localized blurring, distortion and ringing artifacts remained at cortex boundaries close to the skull or air cavities, most prominently in orbitofrontal regions, and in the temporal lobe, above the ear canals, as well as in more inferior slices, particularly in the brainstem. Consequently, such regions might exhibit less sensitive and spatially less defined activation patterns than the ones in visual cortex evaluated here.

We did not evaluate whether our particular choice of B_0_ map resolution or processing contributed to the accomplished image quality or its limitations. Alternative methods to determine B_0_ may provide similar results at reduced scan time, for example, slightly varying TEs during a spiral image time series to estimate the B_0_ map from their phase differences directly (Glover and Law, 2001; Singh et al., 2018) or joint estimation of B_0_ and image from the spiral data itself (Fessler, 2010; Hernando et al., 2008; Patzig et al., 2020). These methods also allow to regularly update B_0_ maps during long fMRI runs, increasing the alignment to the spiral acquisition geometry in case of subject motion.

#### 4.3.4 Iterative Parallel Imaging Reconstruction

To enable single-shot imaging for maximum acquisition efficiency, we also relied on the coil sensitivity profiles for spatial encoding, i.e., parallel imaging. Spiral trajectories are particularly suited for this form of acceleration, because they possess favorable behavior in terms of spatial noise amplification by the coil geometry factor, allowing for higher k-space undersampling (Larkman, 2007; Lee et al., 2021).

For our data with in-plane acceleration factors of 4 using a 32-channel receive array at 7 T this was confirmed through the SD maps of the time series data, which did not exhibit spatially structured residual aliasing or noise patterns. This also points to the robustness of the reconstruction to motion-induced mismatch of measured and actual coil sensitivities.

Compared to Nyquist-sampled spiral data, which could be reconstructed via gridding and conjugate phase correction for static off-resonance effects (Singh et al., 2018 and references therein), parallel imaging of non-Cartesian trajectories typically necessitates iterative reconstruction schemes (Heidemann et al., 2006; Lustig and Pauly, 2010; Pruessmann et al., 2001; Weiger et al., 2002; Wright et al., 2014). Depending on the number of iterations and precision of the off-resonance correction, these algorithms are one or two orders of magnitude slower than direct reconstruction methods. Note, however, that the reconstruction times reported here will not present a general hurdle for deployment, because our Matlab code was not optimized for speed. The numerous matrix-vector multiplications (eq. 5) burden the CG algorithm most, and an implementation on graphical processing units (GPUs) will significantly accelerate reconstruction. High-performance implementations of the conjugate gradient iterative reconstruction algorithm, including off-resonance correction, are publicly available in different MR reconstruction packages, and we have successfully tested reconstruction of the example data presented here in MRIReco.jl (Knopp and Grosser, 2021), written in the modern scientific programming language Julia (Bezanson et al., 2017).

#### 4.3.5 Ultra-high field Magnet (7T) and Gradients

Finally, the availability of an ultra-high field system may be seen as a limitation for the advances presented here. We implemented the spiral sequences at 7T, which has shown particular utility for high-resolution functional MRI due to its superlinear increase in BOLD CNR (Uludağ and Blinder, 2018). From an image reconstruction perspective, this is a challenging scenario, because both static and dynamic field perturbations are exacerbated at ultra-high field and deteriorate conditioning of the expanded signal model. Thus, the adoption of the presented advances in spiral fMRI to lower field strengths (e.g., 3T) not only seems straightforward and worthwhile, but also offers benefits. For example, spiral readouts could be prolonged in light of the more benign field perturbations, mitigating the lower CNR while maintaining high image quality.

Our gradient system, on the other hand, had standard specifications, available on most sites (utilized gradient amplitude 31 mT/m, slew rate 160 T/m/s). Already here, spiral trajectories offered reduced readout times of 19 % compared to EPI due to their higher average speed covering k-space. Because the last 80 % (45 ms) of our high-resolution spiral gradient waveform were amplitude-limited (Fig. 1, black waveform), this acceleration could be considerably increased by dedicated gradient hardware with higher maximum gradient strength, e.g., high-performance whole-body “connectome” gradient coils (Kimmlingen et al., 2012) or insert gradients for head imaging (Foo et al., 2018; Weiger et al., 2018).

### 4.4 Translation to other fMRI applications

This work focused on two-dimensional, slice-selective spiral BOLD imaging. Simultaneous multi-slice (SMS) or 3D excitation schemes offer a complementary means of acceleration, by extending sensitivity encoding to the third encoding (slice) dimension, as, e.g., in stack-of-spiral trajectories (Deng et al., 2016; Engel et al., 2021; Zahneisen et al., 2014), which also provides SNR benefits (Poser et al., 2010). The expanded signal model and image reconstruction framework employed here, apart from the 2D-specific simplifications, are equally applicable to this scenario (Engel et al., 2021; Pruessmann et al., 2001; Zahneisen et al., 2015).

Furthermore, the successful deployment of the in/out spirals here suggests the feasibility of other dual-echo variants, such as out-out or in-in acquisition schemes. In particular, recent correction methods for physiologically or motion-induced noise that rest on multi-echo acquisition (Kundu et al., 2012; Power et al., 2018) could profit considerably from spiral-out readouts: compared to EPI, the shorter minimum TE provides first-echo images with reduced T_2_*- weighting and should enhance disentanglement of BOLD- and non-BOLD related signal fluctuations.

Beyond BOLD, the adaptation of single-shot spiral acquisition for other time series readouts seems promising. In particular fMRI modalities with different contrast preparation (Huber et al., 2017b), such as blood-flow sensitive ASL (Detre et al., 2012, 1992), and blood-volume sensitive VASO (Huber et al., 2018; Lu et al., 2013, 2003) benefit from the shorter TEs offered by spiral-out readouts. These sequences do not rely on T_2_* decay for functional sensitivity, and thus minimizing TE leads to considerable CNR gains (Cavusoglu et al., 2017; Chang et al., 2017).

## Supporting information

Supplementary Material 1: Figure Collection

Supplementary Material 2: Figure Collection

Supplementary Material 4: Slideshow

## Acknowledgments

This work was supported by the NCCR “Neural Plasticity and Repair” at ETH Zurich and the University of Zurich (KPP, KES), the Clinical Research Priority Program of the University of Zurich, CRPP “Pain” (MMS, KES), the René and Susanne Braginsky Foundation (KES), the University of Zurich (KES) and the Oxford-Brain@McGill-ZNZ Partnership in the Neurosciences (NNG, OMZPN/2015/1/3). Technical support from Philips Healthcare, Best, The Netherlands, is gratefully acknowledged. The Wellcome Centre for Integrative Neuroimaging is supported by core funding from the Wellcome Trust (203139/Z/16/Z).

We thank the reviewers for very constructive feedback that helped to considerably improve our manuscript, in particular the concrete suggestions regarding quantification of spatial accuracy of our spiral data via contour distance mapping. We are grateful to Roger Luechinger for technical support with scanning and data curation.

## Conflicts of Interest

At the time of submission, Christoph Barmet and Bertram J. Wilm are employees of Skope Magnetic Resonance Technologies. Klaas P. Pruessmann holds a research agreement with and receives research support from Philips. He is a shareholder of Gyrotools LLC.

## Supplementary Material

**Figure SM 1.** Figure collection of high-resolution (0.8 mm) spiral fMRI data. For every subject, 8 figures are shown, each depicting all 36 slices of the acquisition (top left = most inferior slice, bottom right = most superior slice). All underlay images are based on the functional time series after realignment, and the overlaid t-map was computed from the preprocessed data which included smoothing as well (FWHM 0.8 mm). Order of the figures (1) First volume of spiral fMRI time series; (2) Time-series magnitude mean of spiral fMRI run; (3) Bias-field corrected version of (2); (4) Signal-to-Noise Fluctuation Ratio (SFNR) image of spiral fMRI run (display range 0-30); (5) Standard deviation (SD) image over time of the same run (display range 0-0.05); (6) Coefficient of Variation (CoV) image of the same run, inverse of (4) (display range 0-0.2); (7) Magnitude-mean spiral fMRI image, overlaid with edges of anatomical reference (mean multi-echo spin warp); (8) Overlay of differential t-contrast maps (p < 0.001 uncorrected) on transverse slices of mean spiral image (hot colormap: URLL-ULLR, cool colormap: ULLR-URLL, display range t=3.2-8).

**Figure SM 2.** Figure collection of spiral in/out fMRI data (1.5 mm resolution). For every subject, 24=3×8 figures are shown, i.e., 8 per set of spiral-in, spiral-out, and combined in/out images. Each figure depicts all 36 slices of the acquisition (top left = most inferior slice, bottom right = most superior slice). All underlay images are based on the functional time series after realignment, and the overlaid t-map was computed from the preprocessed data which included smoothing as well (FWHM 0.8 mm). Order of the figures: (1) First volume of spiral fMRI time series; (2) Time-series magnitude mean of spiral fMRI run; (3) Bias-field corrected version of (2); (4) Signal-to-Noise Fluctuation Ratio (SFNR) image of spiral fMRI run (display range 0-30); (5) Standard deviation (SD) image over time of the same run (display range 0-0.05); (6) Coefficient of Variation (CoV) image of the same run, inverse of (4) (display range 0-0.2); (7) Magnitude-mean spiral fMRI image, overlaid with edges of anatomical reference (mean multi-echo spin warp); (8) Overlay of differential t-contrast maps (p < 0.001 uncorrected) on transverse slices of mean spiral image (hot colormap: URLL-ULLR, cool colormap: ULLR-URLL, display range t=3.2-8).

**Figure SM 3.**
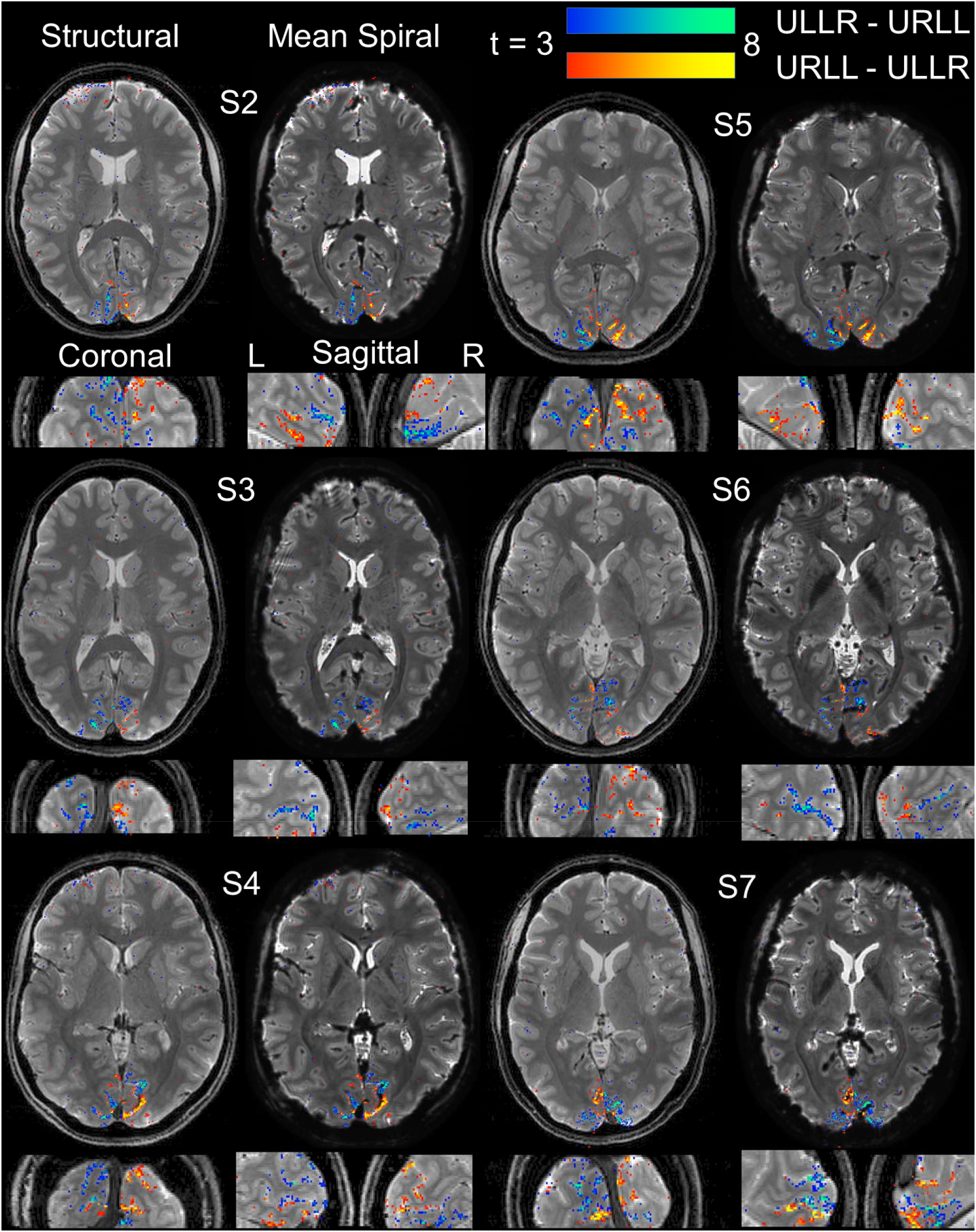
Mean spiral images and activation maps over subjects (S2-S7) for high-resolution spiral-out fMRI, based on unsmoothed data (See Fig. 7 for same visualization using smoothed data). For each subject, the following 4 sections are displayed, with the mean ME image as anatomical underlay: transverse, coronal and sagittal slice (for left (L) and right (R) hemisphere), each chosen for the maximum number of activated voxels (over both differential statistical t-contrasts, p < 0.001 uncorrected). To assess raw spiral data quality, the corresponding mean functional image is displayed side-by-side to the anatomical transverse slice as an alternative underlay.

**Figure SM 4.** Impact of readout duration and spatial smoothing on activation maps in high-resolution spiral-out fMRI data. The animated slideshow toggles between t-maps of the differential contrast (+/− ULLR-URLL), overlaid on the structural image (mean ME) in a single subject (S2): (1) T-map based on smoothed data (FWHM 0.8 mm), thresholded at t > 3.2, (p < 0.001 uncorrected), as presented throughout the other figures, (2) T-map based on unsmoothed data, same thresholding (t > 3.2), and (3) T-map based on unsmoothed data, with more liberal thresholding (t > 2.4, p < 0.01 uncorrected). (4-6) as (1-3), but with spiral-out fMRI k-space data retrospectively cropped to 1 mm resolution before image reconstruction. (1-3) illustrate that moderate spatial smoothing elevates overall CNR (higher t-values) and increases the spatial extent of existing activation clusters in the unsmoothed data. The activation clusters of the smoothed data resemble those in the unsmoothed data at lower thresholds, but without the single-voxel false-positives. This suggests that the averaging of thermal noise via smoothing delivers an implicit cluster size correction. (4-6) illustrate that cropping the spiral readout to 1 mm resolution yields similar activation maps to using the full 0.8 mm readout, both before and after smoothing. This indicates that the assessment of spatial specificity is SNR-limited for the short functional runs (5.5 min) investigated here.

**Table SM 5.**
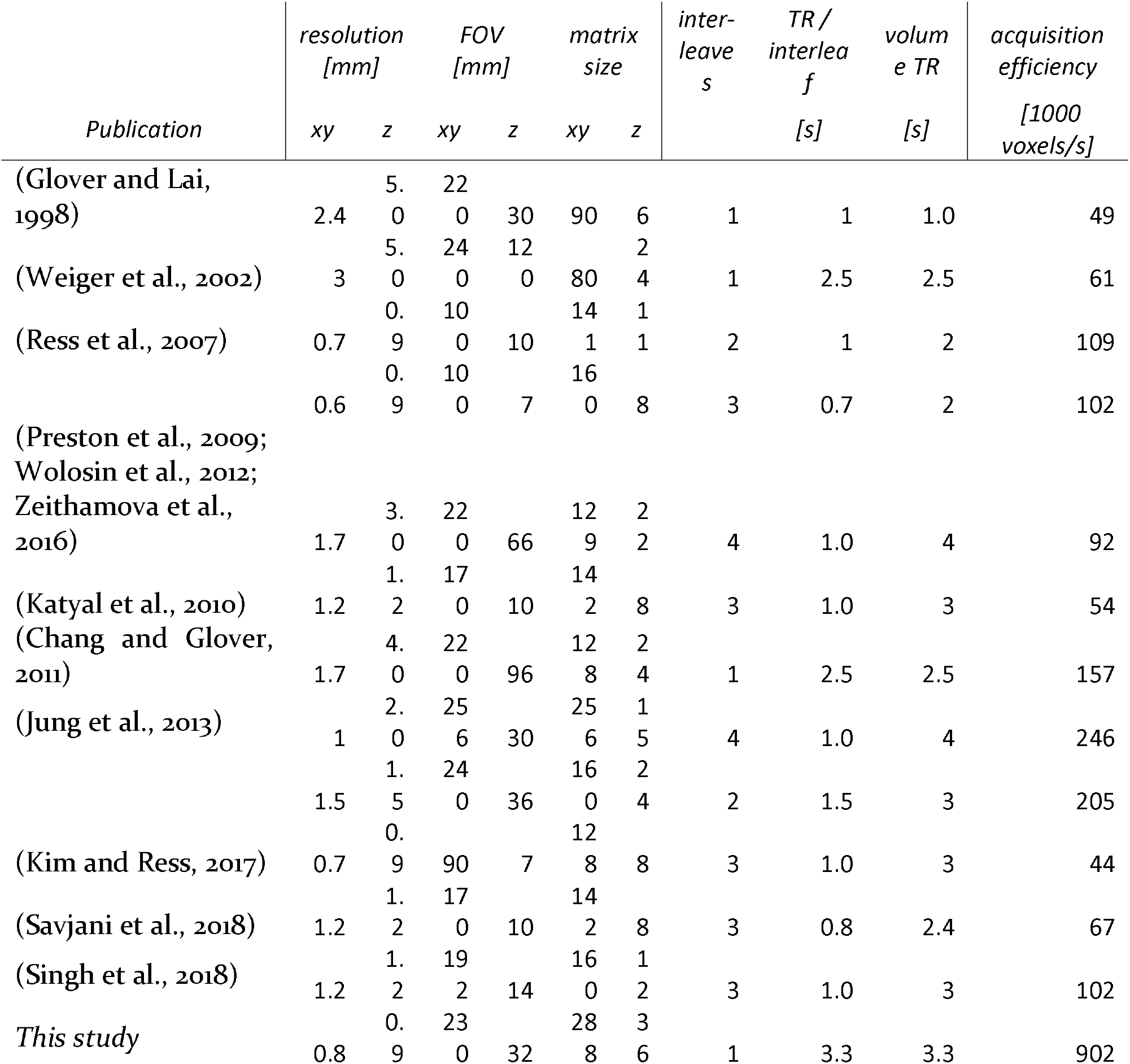
Comparison of acquisition efficiency in published 2D spiral fMRI studies. Nominal acquisition efficiency, computed as resolved voxels per unit time (i.e., matrix size per volume TR) is compared for several publications referenced in this manuscript. The combination of single-shot sequences with long readouts and parallel imaging, as utilized in this study, achieves the highest acquisition efficiency.

**Figure SM 6.**
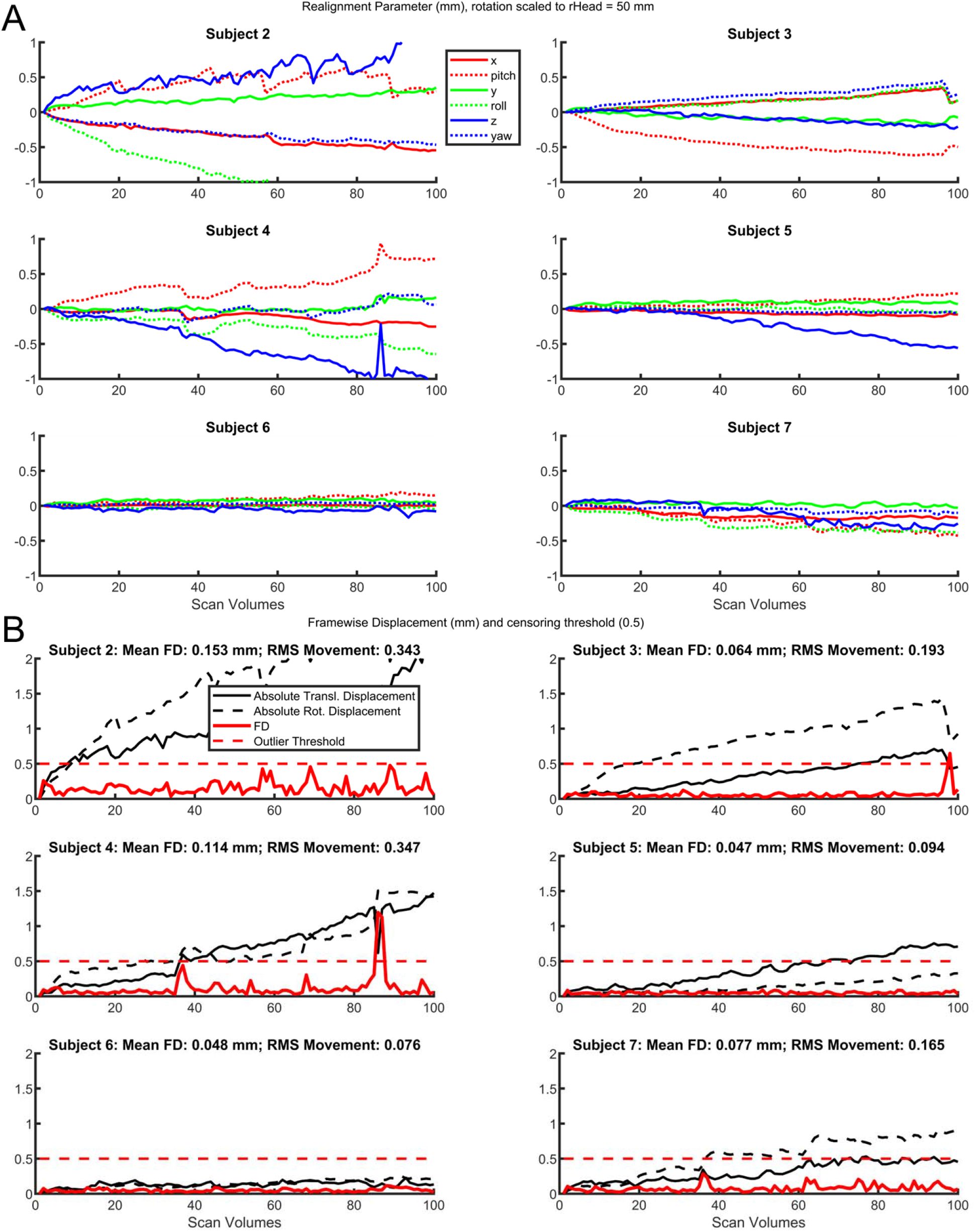
Motion parameters for all subjects during high-resolution (0.8 mm) spiral-out fMRI runs. Displayed are (A) the 3 translational and 3 rotational realignment parameter traces over volumes obtained by rigid-body registration, as well as (B) corresponding framewise displacement (FD) curves (Power et al., 2012). Mean FD was well below 0.2 mm for every subject (max. 0.15 mm, mean FD and standard deviation over subjects 0.09±0.04 mm), which is often considered a very rigorous criterion for censoring of motion-contaminated data (Power et al., 2015). Individual FDs of larger than 0.5 mm occurred in only 3 volumes of 2 subjects (red dashed line threshold).

## Notes

https://github.com/mrikasper/paper-advances-in-spiral-fmri

https://github.com/mrtm-zurich/rrsg-arbitrary-sense

https://dx.doi.org/10.24433/CO.5840424.v1

https://neurovault.org/collections/6086/

https://github.com/mrikasper/julia-recon-advances-in-spiral-fmri

https://doi.org/10.3929/ethz-b-000487412

### Summary of Updates

This version includes changes following the reviewers' comments (2 revision rounds) after submission to the journal NeuroImage. This version is equivalent to the one accepted by the journal.

