## Supplementary figures and images for "Advances in Spiral fMRI: A High-resolution Study with Single-shot Acquisition"

### Supplementary Material 3

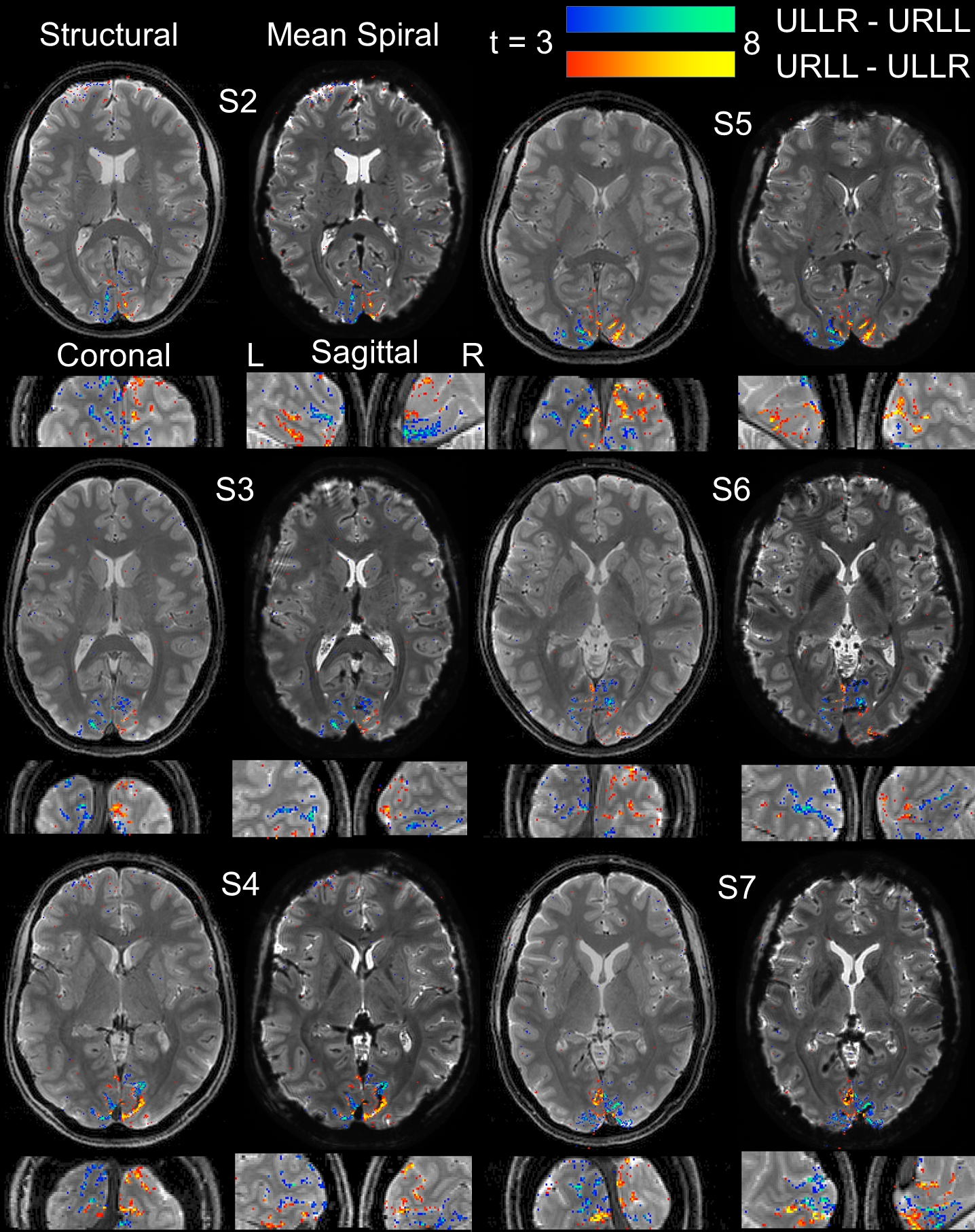

### Supplementary Material 6

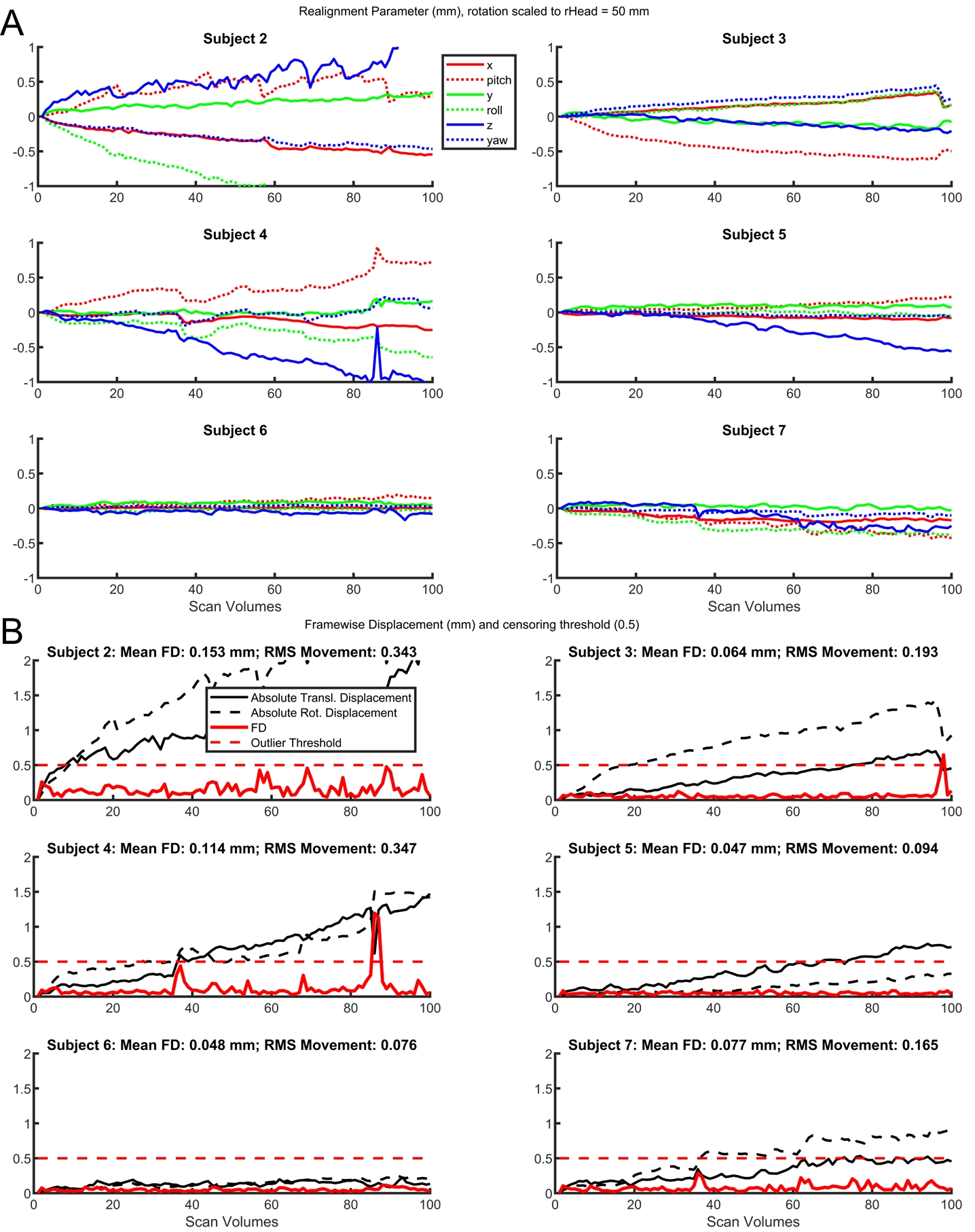
